# Autumn warming decouples the molecular control of photosynthesis and cold acclimation from growth cessation in white spruce

**DOI:** 10.64898/2026.11.23.768595

**Authors:** Bridget K. Murphy, Noelle Perkins, Vera M.E. Velasco, Nathalie Isabel, Jill Wegrzyn, Ingo Ensminger

## Abstract

In evergreen conifers, late-season photosynthetic downregulation is typically coordinated with growth cessation and the induction of cold acclimation, a key component of winter adaptation. However, the transcriptional basis of this coordination and its sensitivity to warming and drought remain poorly understood.
To investigate the molecular basis of these responses, we examined how summer drought, autumn warming and their combination affect the expression of photosynthetic and cold-acclimation genes in white spruce, and whether responses differ among families from different latitudes.
Autumn warming sustained photosynthetic gene expression into early winter while suppressing cold-acclimation genes, demonstrating that photosynthetic and cold-acclimation networks become decoupled from growth cessation, which remained largely unchanged. Although cold acclimation models derived from herbaceous species predict regulation through CBF pathways, canonical CBF/DREB regulators showed little transcriptional response. Instead, warming-induced suppression of cold acclimation was mediated by MYB- and AP2/ERF-centered regulatory networks, indicating a distinct conifer mechanism. Northern and southern families also differed in their transcriptional responses to warming despite exhibiting similar growth-cessation phenology.
Together, these findings identify regulatory networks through which warmer autumns disrupt seasonal coordination in boreal conifers, potentially increasing susceptibility to freezing injury, limiting spring recovery, and ultimately affecting forest carbon sequestration.

**Plain language summary:** Conifers time winter preparation using two cues, shortening photoperiod and declining temperature. Autumn warming delays the temperature cue but not photoperiod. In white spruce, warming sustained photosynthetic gene expression into early winter and suppressed cold-acclimation genes, while growth cessation remained unchanged. Contrary to models from herbaceous plants, this response was not strongly associated with CBF/DREB regulators but with MYB- and AP2/ERF-centered networks.

## Introduction

The magnitude of the terrestrial carbon sink remains uncertain in part because the coupling between carbon uptake and its allocation to woody biomass is poorly understood (Cabon *et al*., 2026). Autumn warming can extend photosynthetic activity in evergreen conifers, but its decoupling from photoperiod-regulated growth cessation may limit the benefits of longer carbon uptake periods for forest growth and carbon sequestration. Recent work from our lab demonstrated this decoupling in white spruce (*Picea glauca)*: experimental warming delayed photosynthetic downregulation and altered latewood xylem development without changing the timing of secondary growth cessation (Fig. S1; see Murphy *et al*., 2025 for all measured traits and methods). These responses also differed between latitudinally distinct families, with the southern family maintaining greater photosynthetic activity under warming than the northern family. Recent evidence from temperate deciduous trees and conifers across broad climatic gradients similarly indicates that extended carbon assimilation and longer growing seasons do not necessarily enhance woody growth (Rao *et al*., 2026; Cabon *et al*., 2026). However, the mechanisms by which warming uncouples photosynthetic activity from growth remain unknown.

Seasonal coordination is central to autumn cold acclimation in evergreen conifers. Growth cessation is strongly regulated by photoperiod in many evergreen species (Rossi *et al*., 2006; Hamilton *et al*., 2016; Wallin *et al*., 2017), whereas photosynthetic downregulation, cold hardening, and xylem differentiation are highly responsive to temperature (Begum *et al*., 2012; Chang *et al*., 2016; Stinziano & Way, 2017). After cambial cell production ceases, recently produced xylem cells continue secondary-wall thickening and lignification, leaving latewood development sensitive to autumn temperature and carbon availability (Rathgeber *et al*., 2016; Cartenì *et al*., 2018; Pampuch *et al*., 2021). Declining autumn temperatures coordinate reductions in growth-driven carbon demand and photosynthetic activity with increases in carbon storage, photoprotection, and freezing tolerance (reviewed in Chang *et al*., 2021). However, warming may delay these temperature-sensitive adjustments without altering photoperiod-regulated growth cessation, creating a mismatch among processes that normally occur in concert during cold acclimation (Busch *et al*., 2007, 2008). Additionally, summer drought may influence autumn acclimation indirectly through carry-over effects on carbon balance, hydraulic status, and stress signalling, potentially modifying subsequent responses to autumn warming (D’Orangeville *et al*., 2018; Wang *et al*., 2024).

These coordinated seasonal adjustments are underpinned by extensive transcriptome reprogramming across photosynthetic, developmental, and stress-response pathways. In Norway spruce (*Picea abies*), seasonal transcriptome profiling identified a distinct autumnal acclimation phase characterized by downregulation of plastid RNA machinery and upregulation of cold-responsive gene categories (Bag *et al*., 2021). Although the ICE–CBF–COR signaling cascade is a central regulator of cold acclimation in *Arabidopsis thaliana* (Stockinger *et al*., 1997; Fowler & Thomashow, 2002; Cook *et al*., 2004; Park *et al*., 2015), its role in conifers remains uncertain. Conifers express CBF-like homologues, COR genes such as dehydrins, and other cold-responsive genes involved in soluble sugar accumulation, membrane stabilization, and hormonal signaling. However, evidence increasingly suggests that cold acclimation in evergreens involves substantial CBF-independent regulation, including members of the WRKY, NAC, and AP2/ERF transcription factor families (Holliday *et al*., 2008; Vergara *et al*., 2022; Rathore *et al*., 2022).

Local adaptation may generate population-level differences in the timing and temperature sensitivity of autumn transcriptome reprogramming. Provenances vary in the photoperiodic thresholds that regulate growth cessation and dormancy induction, with northern provenances typically initiating cold acclimation and downregulating photosynthesis earlier than southern provenances (Holliday *et al*., 2008; Sebastian-Azcona *et al*., 2018; Fréchette *et al*., 2020). This genetic differentiation may also extend to the regulation of CBF-dependent and CBF-independent pathways, but whether it shapes transcriptional responses to autumn warming and summer drought remains unknown. White spruce is well suited to investigating these differences because it exhibits substantial genetic variation in phenological, structural, and physiological traits across its range (Beaulieu *et al*., 2011; Sebastian-Azcona *et al*., 2018; Depardieu *et al*., 2020; Leites & Benito Garzón, 2023). White spruce is also one of the most widely planted and ecologically important boreal tree species in Canada, playing a central role in forest productivity and reforestation (Mullin *et al*., 2011).

The contrasting photosynthetic responses previously observed in northern and southern white spruce families provided an opportunity to investigate the molecular basis of warming-induced decoupling between photosynthetic activity and autumn acclimation. To overcome the challenges of complex conifer genomes, the long-read transcriptome assembly generated in Murphy et al. (in press) was used to supplement the existing white spruce annotation (Gagalova *et al*., 2022). We asked whether autumn warming disrupts the seasonal coordination of transcriptional networks associated with photosynthesis, xylem development, photoprotection, and cold acclimation, and whether these responses are modified by prior summer drought or family background. We predicted that warming would sustain photosynthetic and xylem-related transcriptional programs into late autumn while delaying or suppressing cold acclimation pathways, resulting in progressive decoupling of processes that are normally synchronized during the seasonal transition to winter. We further predicted that the southern family would exhibit a later autumn transcriptional transition than the northern family, consistent with its greater maintenance of photosynthetic activity under warming. Finally, we evaluated whether these responses were associated with canonical CBF/DREB-mediated regulation or alternative transcription factor networks. To address these questions, we combined contrast-based differential expression, gene-set enrichment, candidate gene analyses, and co-expression network approaches spanning summer through early winter (Fig. 1).

**Figure 1.**
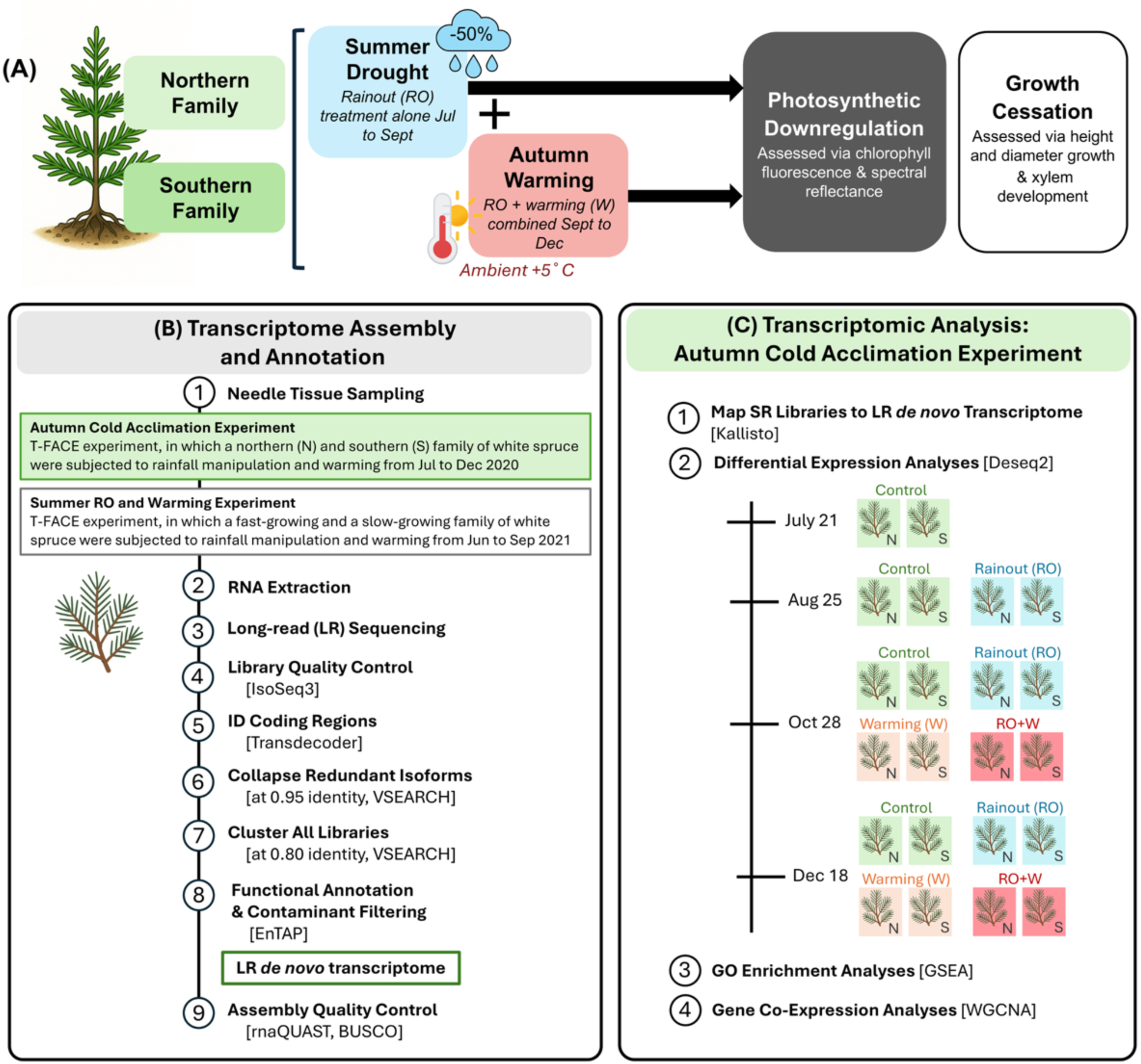
Bioinformatics pipeline used in this study. (A) Schematic of the experimental design: seedlings from northern and southern families were exposed to summer rainout (RO), then autumn warming (W), either alone or in combination (RO+W). (B) Workflow for generating the LR *de novo* transcriptome assembly from needle tissue sampled across the Autumn Cold Acclimation and Summer RO and Warming experiments. (C) Workflow for gene expression analyses for the Autumn Cold Acclimation Experiment, including mapping of short-read (SR) libraries to the LR assembly and downstream differential expression and enrichment analyses. Software used at each step is shown in brackets. N, northern family; S, southern family.

## Materials and Methods

### Plant Material and Experimental Design

Transcriptomic analyses used RNA-seq data from a field-based T-FACE experiment at the Koffler Scientific Reserve, King City, Ontario, Canada (44°01′44.3″ N, 79°32′19.2″ W). Two white spruce (*Picea glauca*) families representing northern and southern origins were sampled across four climate treatments: control, rainout (RO), warming (W), and combined RO+W (Fig. 1A; Murphy *et al*. 2025). The first family was generated from a controlled cross between a northern parent from Lac Simard, Québec and a more-southern parent from Petawawa, Ontario, and the second family was generated from a cross between two southern parents from Hungerford, Ontario. Seedlings were produced by Natural Resources Canada (Laurentian Forestry Centre) in 2017, and parental origins were chosen to capture variation in temperature and photoperiod sensitivity along a latitudinal gradient of ∼400 km.

The field experiment followed a step-wise, factorial manipulation of reduced soil water content (rainout, RO) and warming implemented in a T-FACE design at the Koffler Scientific Reserve (King City, Ontario, Canada). Four treatments were applied: control, RO, warming (W), and combined RO+W. Treatments were replicated in three plots each. Rainout structures were installed on July 22 and maintained until November 27, and warming began on August 31 and continued until the end of the experiment. Warming was applied using infrared heaters that increased leaf temperature by approximately 5 °C during both day and night (see Fréchette *et al*., 2020 for T-FACE technical details; Murphy *et a*l. 2025 for full site implementation). Rainout structures intercepted approximately 50% of ambient precipitation, and non-RO plots contained dummy profiles to control for structural artefacts. Plot-level air temperature and humidity, soil temperature and moisture, and infrared leaf temperature were logged at 30-min intervals, and an on-site weather station recorded meteorological variables at 15-min intervals (Fig. S1A; Method S2). Mature needles were sampled from four-year-old seedlings at 16:00 h in late July, August, October, and December, flash-frozen in liquid nitrogen, and stored at −80 °C until RNA extraction.

### RNA Extraction and Sequencing

Total RNA was extracted from 150 mg of homogenized needle tissue using a cetyltrimethylammonium bromide-based protocol adapted from Chang *et al*.(Chang *et al*., 1993), followed by purification with the RNeasy Mini Kit and on-column DNase digestion (Qiagen, Germany). RNA concentration and integrity were assessed using a Qubit 3.0 fluorometer and an Agilent 2100 Bioanalyzer with an RNA Nano 6000 chip. All samples had RNA integrity numbers exceeding 6.7. Samples from three seedlings per family, treatment, and sampling date were sequenced, yielding 66 RNA-Seq libraries. Libraries were prepared using a stranded mRNA protocol (NEB) and sequenced at Génome Québec (Montreal, QC, Canada) on an Illumina NovaSeq 6000 platform with S4 chemistry to generate 100 bp paired-end reads. Libraries were multiplexed at 1/22 per lane.

### Short-read Processing and Mapping

Raw reads were quality-filtered using Trimmomatic v0.36 to remove adapter sequences, low-quality bases, and reads shorter than 30 bp (Bolger *et al*., 2014). Read quality was assessed using FastQC v0.11.7 (Andrews, 2010). High-quality reads were pseudo-aligned with Kallisto v0.44.0 (Bray *et al*., 2016) to a long-read *de novo* white spruce transcriptome assembled from needle tissue using PacBio Iso-Seq (Murphy *et al*. in press), which served as the reference for all downstream expression and annotation-based analyses (Fig. 1B). Twenty-two of the 24 samples used for long-read transcriptome assembly were also included in the short-read dataset, ensuring concordance between sequencing platforms. Details of transcriptome assembly, quality control, and functional annotation are provided in Murphy *et al*. (in press) and Methods S1.

### Differential Expression Analysis

Transcript abundance estimates generated with Kallisto v0.44.0 were analyzed using DESeq2 (Love *et al*., 2014). Estimated counts were normalized using DESeq2’s internal normalization procedure, and log₂ fold changes (LFCs) were estimated and shrunken using ‘normal’ to improve effect-size stability. Differentially expressed transcripts (DETs) were identified using an absolute LCF threshold of ≥ 1 and a Benjamini–Hochberg–adjusted P-value (false discovery rate, FDR) cutoff of 0.05.

Analyses were conducted separately for each sampling date using targeted pairwise contrasts. To assess family-specific responses to treatments, family and treatment were combined into a single grouping factor, allowing contrasts both within families across treatments and between families within treatments. Seasonal transcriptomic changes associated with autumn cold acclimation were evaluated under control conditions by comparing July to August, October, and December within each family. Global expression patterns were examined using variance-stabilized counts and visualized through principal component analysis, MA plots, and volcano plots generated with ggplot2 (Wickham, 2016). One RO+W sample from December consistently clustered with the opposite family, indicating a labeling error, and was excluded from all subsequent analyses (Fig. S2).

### Gene Set Enrichment Analysis

Gene set enrichment analyses were performed using a rank-based approach implemented in the fgsea R package v1.32.4 (Korotkevich *et al*., 2021). Transcripts were ranked using DESeq2 Wald statistics for each contrast, and enrichment was assessed using Gene Ontology (GO) term sets derived from functional annotations of the long-read white spruce transcriptome. Enrichment significance was evaluated using normalized enrichment scores (NES) and false discovery rate (FDR) values calculated with the fgseaMultilevel algorithm. For each enriched GO term, fgsea reports a leading-edge subset, defined as the transcripts contributing most strongly to the enrichment signal. This rank-based approach leveraged the full transcriptome to enable detection of coordinated but modest expression shifts across functionally related genes that may not be captured by threshold-based over-representation analyses.

### Identification of Candidate Genes Underlying Warming Responses in Photosynthetic and Xylem Pathways

Putative candidate transcripts were selected to identify genes that likely contributed to the phenotypic responses observed under warming, particularly within photosynthesis- and xylem development-related pathways. Candidates were identified from differentially expressed transcripts (DETs) with functional annotations based on GO and KEGG terms. Transcript annotations represent putative white spruce homologs derived from sequence similarity and gene family assignment. Candidates were required to be differentially expressed in at least two contrasts, spanning at least two combinations of sampling date, family, and treatment. Within each pathway, transcripts were ranked by the number of significant contrasts, with ties resolved using the median absolute log2 fold change. Only the top ten candidates per category per contrast type were retained for visualization to limit complexity.

### Transcription Factor–Specific Analysis

Transcription factor annotations were obtained from the annotated long-read white spruce transcriptome, as described in Murphy *et al*. (in press). TF transcripts were intersected with DESeq2 differential expression results, and differentially expressed TFs were summarized by family for each treatment and sampling date. Heatmaps were generated using variance-stabilized expression values from DESeq2. Expression was centered and scaled by transcript using Z-score transformation prior to hierarchical clustering, enabling visualization of relative seasonal and treatment-associated expression patterns across transcription factor families, treatments, and sampling dates.

### Weighted Gene Co-Expression Network Analysis

Transcript expression estimates generated with Kallisto v0.44.0 (Bray *et al*., 2016) were filtered to remove low-abundance transcripts using edgeR v4.4.2 (Robinson *et al*., 2010). Sample quality was assessed using hierarchical clustering, with no outliers detected. A signed gene co-expression network was constructed using WGCNA v1.73 (Langfelder & Horvath, 2008), applying blockwise module detection with a minimum module size of 30 and a merge cut height of 0.25. A soft-thresholding power of 12 was selected to approximate scale-free topology. Modules were summarized using eigengenes, and network structure was visualized using dendrograms and topological overlap heatmaps.

Gene set enrichment analysis of biological GO terms was performed for all modules using fgsea, with transcripts ranked by module membership. Hub genes were identified as genes with high module membership, defined as an absolute correlation of ≥ 0.80 with the corresponding module eigengene. Seasonal effects were evaluated using a December-only subset by recalculating eigengenes and module correlations while retaining module assignments from the full dataset.

### Identification of Regulatory Genes for Pathway Reconstruction

Genes and gene families included in the regulatory pathway diagram were identified using the cold acclimation framework outlined in Chang *et al*., (2021), with downstream gene families described by Holliday *et al*. (2008). Putative homologs were identified from the long-read *de novo* white spruce transcriptome using similarity search results generated during EnTAP v2.3.0 functional annotation (Hart *et al*., 2020). Similarity searches against the Araport11 and RefSeq plant protein databases were examined in parallel. Transcript identifiers corresponding to each annotated homolog were then cross-referenced with RNA-Seq differential expression results to determine whether annotated homologs were differentially expressed under W, RO, or combined RO+W treatments. Conserved relationships were inferred based on homology and functional annotation, and directionality of regulatory responses was assigned using LFC estimates relative to control conditions.

## Results

### RNA sequencing and library summary

Needle sampling for RNA-Seq was conducted in July, August, October, and December, matching the seasonal physiological measurements reported in the companion field experiment (Murphy *et al*. 2025; Fig. 1C; Fig. S1A). Sequencing of 66 white spruce RNA libraries yielded 9 billion paired-end reads, with 106-168 million reads per library (Table S1). After quality filtering, 104-165 million reads per sample were retained. Across libraries, 64–79% of reads were pseudoaligned to the long-read *de novo* transcriptome, with a mean alignment rate of 75%.

### Seasonal transcriptional changes in photosynthetic, growth, stress, and hormonal pathways during the autumn-to-winter transition

We first characterized baseline seasonal shifts in transcript expression. For this, we identified differentially expressed transcripts (DETs) in pairwise contrasts of July vs August, October, and December within each family under ambient conditions (Table S2). Both families showed extensive transcriptional changes as autumn progressed toward winter. From July to August, the southern family had 136 DETs, whereas the northern family had 1,103 DETs (Fig. 2A). DET counts increased sharply from July to October (southern, 8,248; northern, 8,011) and July from December (southern, 11,221; northern, 10,736). Overlap in DETs between families transiently increased as temperature declined and photoperiod shortened, indicating greater convergence of transcriptional responses under shared environmental conditions (Fig. 2B).

**Figure 2.**
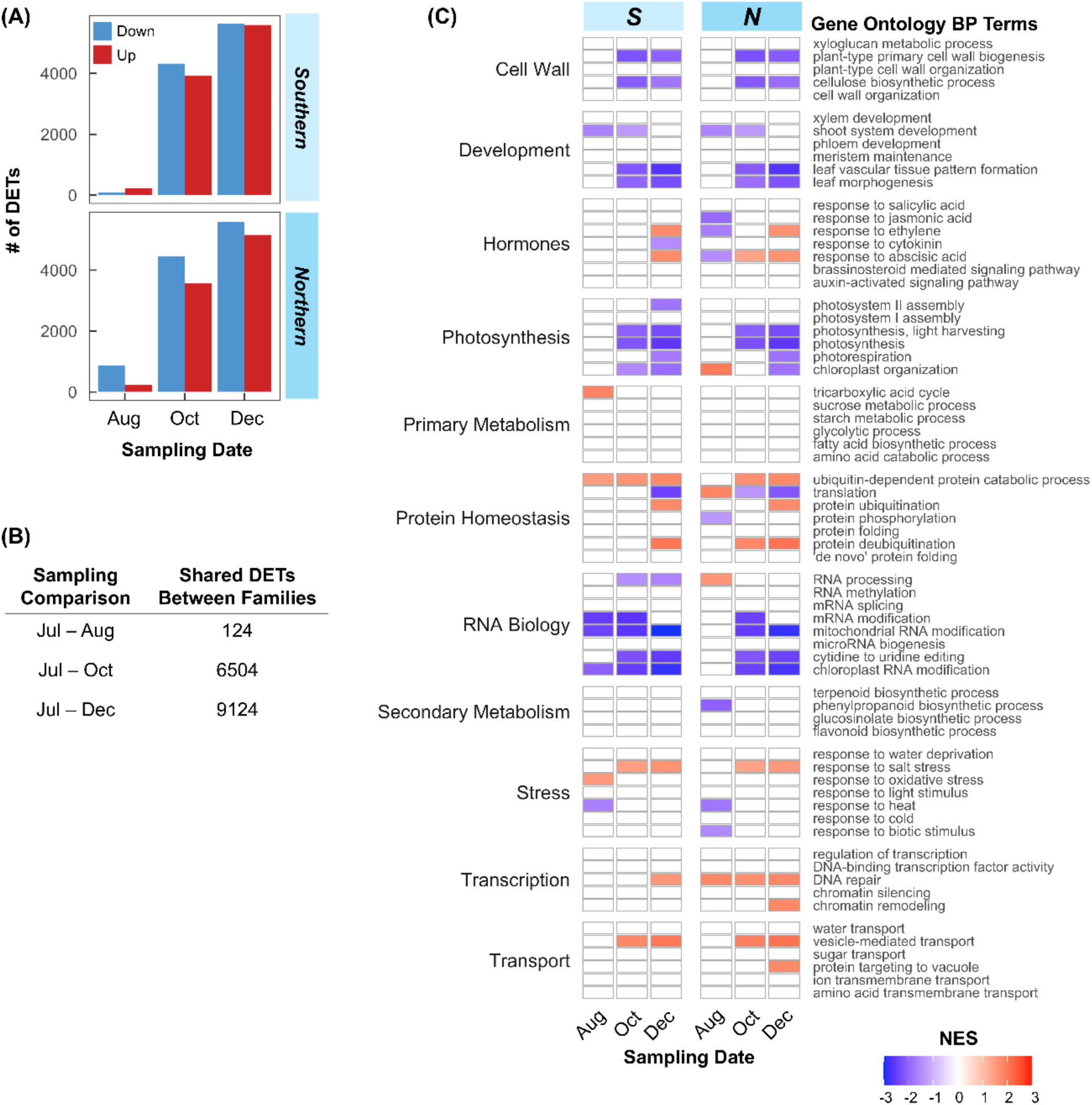
Seasonal shifts in gene expression under ambient conditions relative to early summer. Differentially expressed transcripts (DETs) were identified within each family under control conditions by comparing later sampling dates (August, October, December) to the July time point. (A) Numbers of upregulated (red) and downregulated (blue) DETs for Southern and Northern families. (B) Numbers of DETs shared between families for each July-based comparison. (C) Gene set enrichment analysis (GSEA) of Gene Ontology Biological Process (BP) terms, shown as normalized enrichment scores (NES) for each family and comparison, with positive NES indicating enrichment of upregulated pathways and negative NES indicating enrichment of downregulated pathways.

Gene set enrichment analysis revealed coordinated functional shifts accompanying these seasonal transitions (Fig. 2C; Table S3). From July to October and July to December, gene sets associated with photosynthesis (e.g., ‘photosynthesis, light harvesting’ and ‘chloroplast organization) and growth (e.g., ‘shoot system development’ and ‘plant organ development’) were negatively enriched. Many of the leading-edge genes, which contributed most strongly to these enrichment signals, were also detected as downregulated DETs, including *LHCA2*, *GAPB*, and *GA20Ox. GA20ox is a* gibberellin biosynthesis gene previously linked to short-day-induced growth cessation in poplar (Eriksson *et al*., 2015). By contrast, gene sets related to hormonal signaling (e.g., ‘response to ethylene’ and ‘response to abscisic acid’), stress (e.g., ‘DNA repair’, and ‘response to salt stress’), and protein homeostasis (e.g., ‘protein ubiquitination’) were positively enriched. The most pronounced signals were observed in the July to December comparison. Representative upregulated DETs included ABA signaling genes (e.g., *ABI5-like*), ABA-independent signaling components (e.g., *ERFC3*), and stress-associated proteins (e.g., *COR47-like*).

### Treatment-induced shifts in gene expression across and between families

Transcriptional responses to treatment and between families differed in both magnitude and seasonal timing, with varying overlap. Across treatments and sampling dates, 2,693 unique treatment-responsive DETs and 4,807 family-level DETs were identified (Fig. 3A,B; Table S4). Within the RO treatment, the largest transcriptional response was observed in August, particularly in the Northern family, whereas responses to W and RO+W increased from autumn to early winter (Fig. 3A). By October and December, treatment responses were dominated by transcript downregulation. Family-level contrasts revealed pronounced variation in the number of differentially expressed transcripts across treatments and sampling months (Fig. 3B; Table S4).

**Figure 3.**
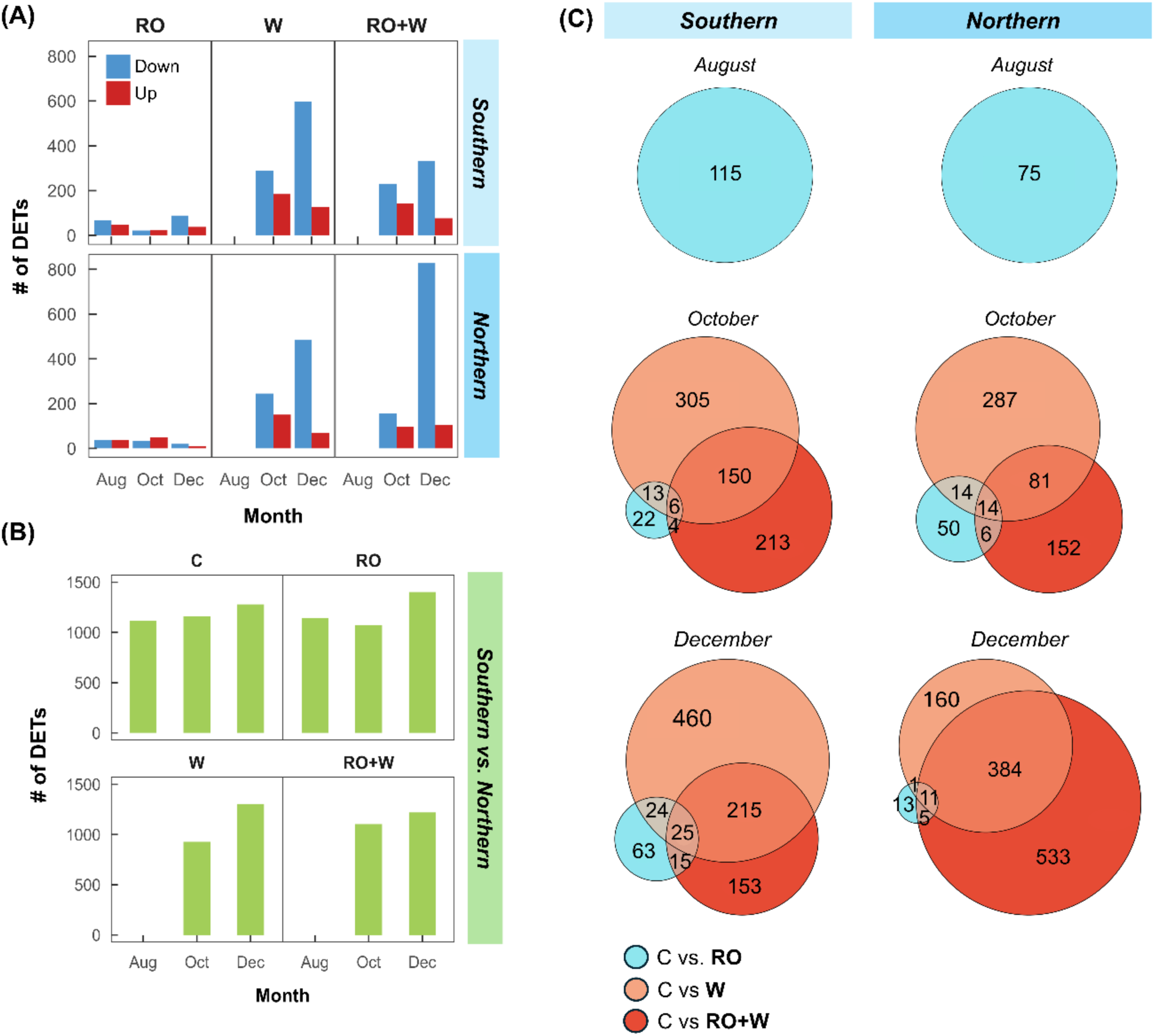
Treatment-induced shifts in differential gene expression within and between northern and southern white spruce families. (A) Numbers of differentially expressed transcripts (DETs) in each family under rainout (RO), warming (W), and combined RO+W, relative to the control (C) at each sampling date. Upregulated DETs are shown in red and downregulated DETs in blue. (B) Numbers of DETs between families (southern vs northern) within each treatment across sampling dates. (C) Area-proportional Venn Diagrams showing overlap of treatment-responsive DETs (C vs RO, C vs W, C vs RO+W) within southern (left) and northern (right) families at each sampling date. “NA” indicates time points when only the rainout treatment was applied. DETs were identified by a |log2 fold change| > 1 and an FDR < 0.05.

Approximately one-quarter (1,369) of these DETs overlapped between treatment-level and family-level contrasts, whereas the remainder were unique to either treatment or genetic background. Overlap among treatment-responsive transcripts was greatest for W and RO+W in December (Fig. 3C; Table S5). In contrast, few transcripts were shared between RO and RO+W (southern: 15; northern: 5) and between RO and W (southern: 24; northern: 1).

### Functional enrichment patterns across treatments and families

Gene set enrichment analyses revealed consistent but seasonally structured functional responses across treatments and families (Fig. S3; Tables S6 and S7). GO terms associated with transcription and plastid organization were frequently enriched, whereas membrane-associated and stress-response GO terms tended be negatively enriched, indicating coordinated downregulation of these pathways. Gene sets directly associated with abiotic stress (e.g., ‘response to cold’ and ‘response to heat’) showed limited enrichment (Fig. 4A). However, the gene set ‘defense response’ was negatively enriched, which may reflect reduced expression of genes involved in cryoprotective responses. Plant antifreeze proteins are homologous to three classes of pathogenesis-related proteins: chitinases, B-1,3-glucanases, and the thaumatin family (Hon *et al*., 1995), which comprised the leading-edge genes contributing to this signal. Gene sets related to signaling and regulatory pathways were also negatively enriched, indicating broad repression of stress-related and cellular response pathways, under W and RO+W in both families. Several of these gene sets also differed between families (Fig. 4B).

**Figure 4.**
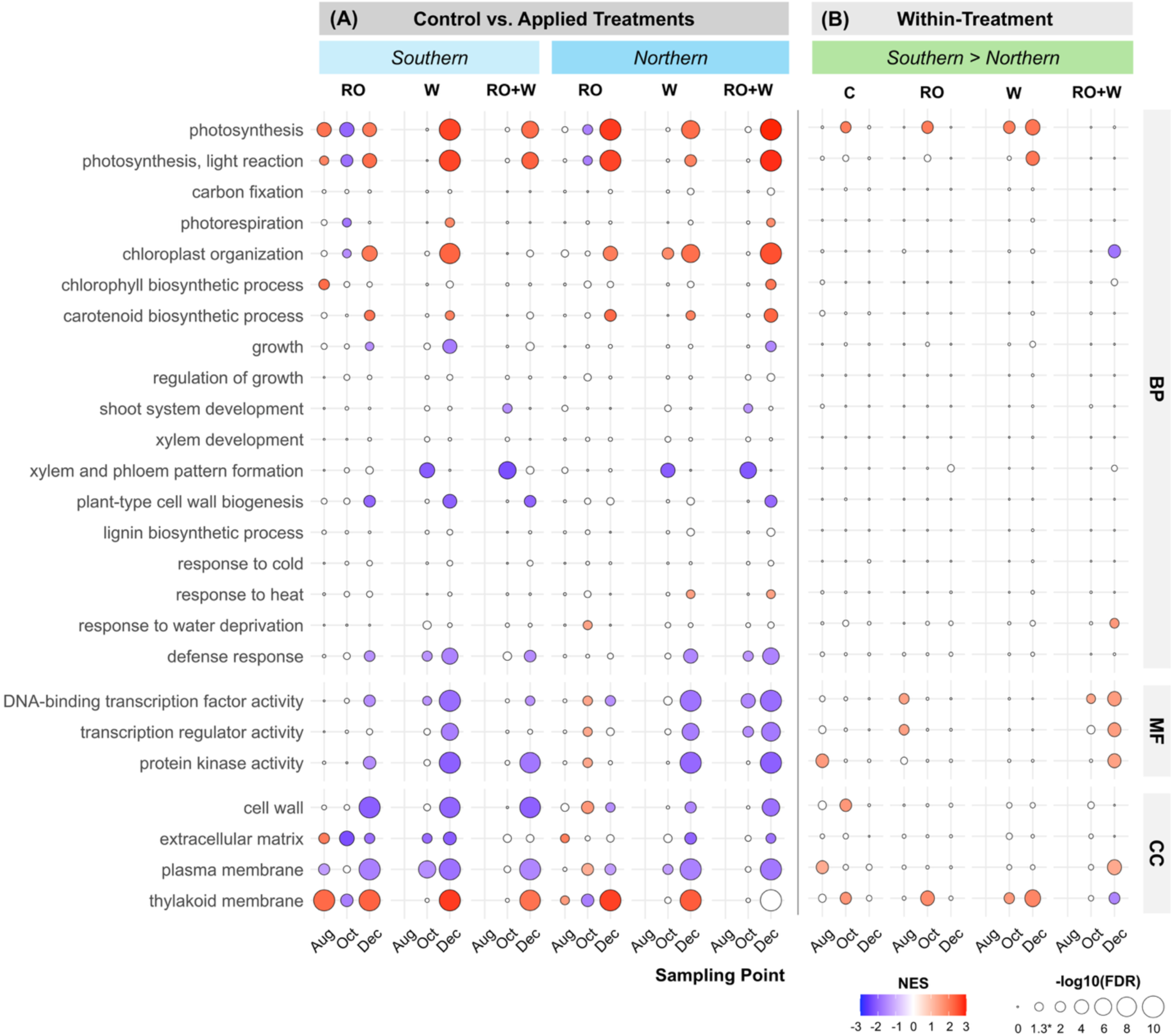
Gene Ontology (GO) enrichment patterns associated with autumn cold acclimation across treatments, families, and sampling dates. Bubble plots show gene set enrichment analysis (GSEA) results for selected GO terms across Biological Process (BP), Molecular Function (MF), and Cellular Component (CC) categories. (A) Control versus applied treatment contrasts (RO, W, RO+W) shown separately for Southern and Northern families. (B) Within-treatment contrasts comparing Southern versus Northern families for each treatment, including the control. All contrasts were performed within sampling dates. Bubble size denotes enrichment significance (−log10[FDR]). Bubble colour indicates normalized enrichment score (NES) for significant terms only (FDR < 0.05), with red indicating positive and blue indicating negative NES. Positive NES reflects enrichment toward transcripts higher in the applied treatment relative to control (A) or higher in the Southern relative to the Northern family (B). C, control; RO, rainout; W, warming; RO+W, combined rainout and warming.

Photosynthesis-, pigment-, and thylakoid-associated gene sets showed repeated positive enrichment under W and RO+W in December (Fig. 4A). Within treatments, the southern family exhibited stronger positive enrichment of photosynthesis-associated processes than the northern family in October and, under warming, in December (Fig. 4B), indicating a sustained transcriptional activation of photosynthetic pathways. Although some photosynthesis-related gene sets were also enriched under RO alone (Fig. 4), this enrichment did not translate into DETs. Enrichment of growth- and xylem-associated processes was less frequent and more variable. Differences in growth- and xylem-associated enrichment between families were not detected within treatments (Fig. 4B). These enrichment patterns are further resolved at the gene level for photosynthesis- and xylem-associated processes in the following section.

### Treatment- and family-dependent expression of candidate transcripts in photosynthesis and xylem-associated pathways

Gene set enrichment analyses identified broad pathway-level shifts, but only a subset of transcripts showed pronounced and repeated differential expression. These transcripts were prioritized to identify putative genes associated with the downstream photosynthetic and xylem-related responses observed under warming. GO and KEGG terms used for pathway assignment are provided in Table S8. Transcript annotations represent putative white spruce homologs derived from sequence similarity and gene family assignment. Accession numbers and reference species are provided in Tables S9 and S10.

Within photosynthesis and associated carbon metabolism pathways, candidate transcripts were enriched for Calvin cycle and chloroplast-associated functions. Representative candidates included enzymes involved in carbon fixation and regeneration, such as *FBPase-like* and *GAPB* (Fig. S4; Table S9). Stress-associated genes linked to redox regulation and protein protection were also identified, including *CAT-like, sHSP21-like*, and *FTSH2.* Under W and RO+W, Calvin cycle-associated genes showed higher expression in October and December relative to the control, whereas stress-associated candidates showed more variable responses (Fig. S4). Additional candidates related to photosynthetic electron transport, photosystem assembly, light harvesting, and photoprotection, including *PGR5-like, PSA2, PPD4, LHCA1,* and *CYP37,* also showed treatment- and family-dependent expression patterns (Fig. S4 and S5; Table S9).

Within xylem- and cell wall-associated pathways, candidate transcripts frequently showed reduced expression under warming (Fig. S5; Table S10). These included genes associated with cellulose synthesis, hemicellulose remodeling (e.g., *CESA5-like, XTH8*) and oxidative cell wall modification (e.g., *PER67-like*). RO alone had minimal effects on most candidates, with the exception of *XTH2*. Family-level differences in xylem-associated transcript abundance were detected but were weaker and less consistent than those observed for photosynthesis-related candidates (Fig. S5). Family contrasts indicated lower expression of transcripts associated with carbohydrate metabolism and sugar signaling (e.g., *TPS1*) and lignin biosynthesis (e.g., *CCOAOMT*) in the southern family (Fig S6). Transcripts associated with oxidative cell wall modification (e.g., *PER, class III*) were higher in the southern family.

### Seasonal and treatment-dependent changes in transcription factor expression

We examined transcription factor (TF) expression across treatments and sampling dates. Responses of TF families were limited in August but expanded markedly in October and December (Fig. S6; Table S11). Most differentially expressed TFs showed reduced transcript abundance under W and RO+W. This pattern was most pronounced within the AP2/ERF and MYB families, followed by WRKY, NAC and other bHLH families (Fig. S6). RO alone elicited relatively few TF responses compared to warming treatments.

Clustering of the 50 TFs with the most variable expression identified two major groups (Fig. 5; Table S12). Cluster 1 showed reduced transcript abundance under W and RO+W in October and included homologs of *A. thaliana* TFs associated with abscisic acid signaling, oxidative stress responses, leaf development, and cold-response regulation, such as *MYB15*, *ERF3, ANAC035, and ANAC02*. Most TFs in this cluster shifted toward control and RO expression levels by December, although *MYB15* and *ANAC035* remained downregulated under W and RO+W. Cluster 2, located at the bottom of the heatmap, exhibited a gradual decline in transcript abundance from August through December across all treatments and included TFs associated with light and photoperiod signaling, such as *bZIP16* and *NF-YC4*. Outside these two clusters, additional TFs related to cold acclimation were upregulated from July to October and from July to December, including *DREB2C*, *WRKY2*, and *PRR2*.

**Figure 5.**
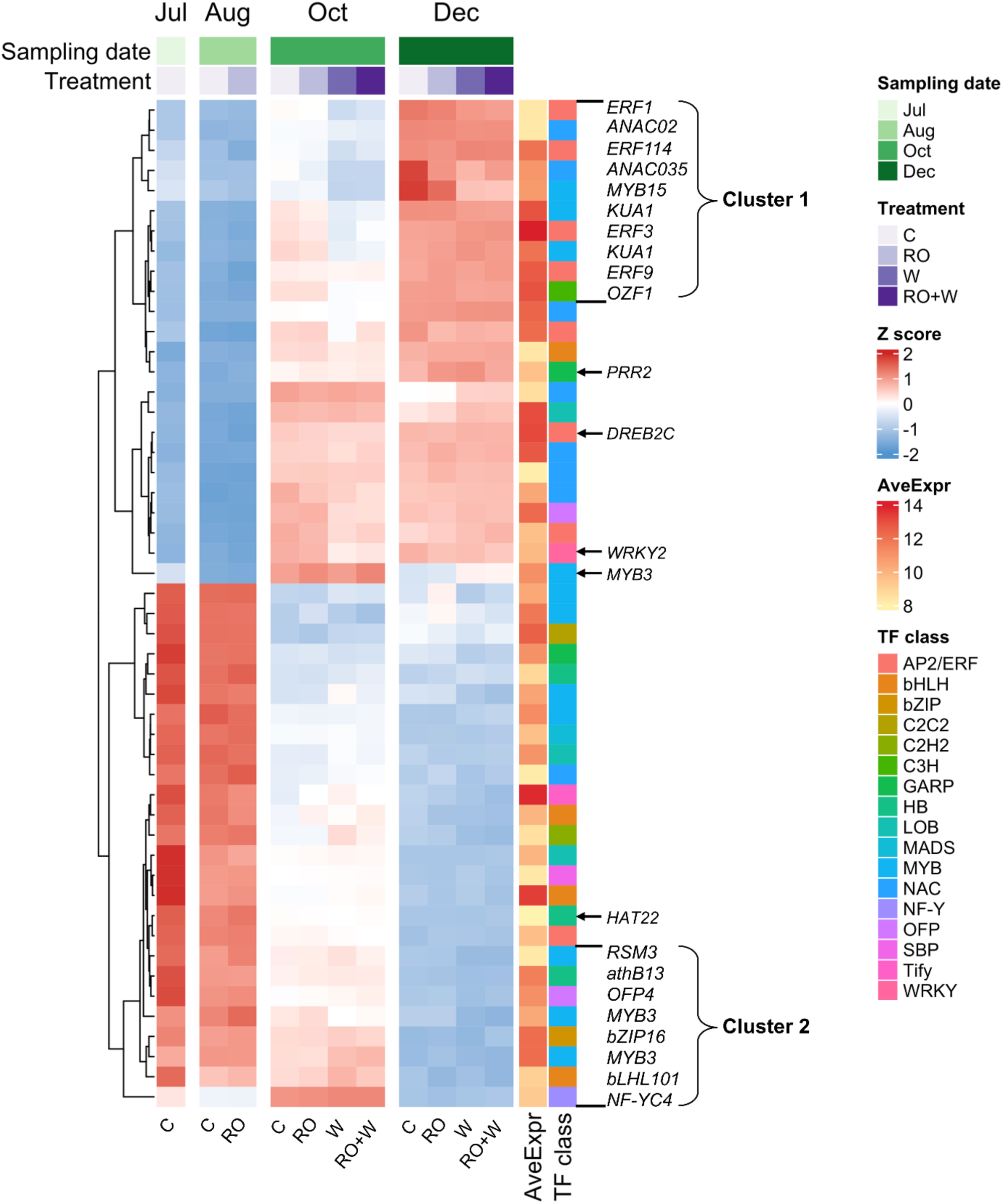
Heatmap showing expression patterns of the 50 most variable transcription factors across sampling months and treatments. Columns correspond to treatments (C, RO, W, RO+W) within each month (July–December), and rows represent individual transcription factor transcripts. Expression is displayed as row-scaled Z-scores, with hierarchical clustering used to identify groups of transcripts exhibiting similar temporal trajectories. Annotation bars indicate sampling month, treatment, average variance-stabilized expression (AveExpr), and transcription factor family.

### Gene co-expression analysis to assess genetic background and treatment-dependent cold acclimation responses

Gene co-expression network analysis was conducted across families, treatments, and sampling dates. The analysis identified 21 modules, with overall network structure driven primarily by genetic background. Family exhibited the most significant and pervasive associations across modules (Fig. S7), indicating substantial genetic control over coordinated gene expression. Treatment effects were most pronounced in late season. To better resolve responses to warming, module-trait relationships were re-examined using December samples, corresponding to the period of maximal warming response. Four modules showed particularly strong associations with either family or treatment (Fig. 6A). Two modules, MEmidnightblue and MEtan, showed treatment-associated downregulation of eigengene expression under W and RO+W compared to C and RO (Fig. 6B). Two additional modules primarily reflected genetic divergence between families. The direction and relative strength of these associations were consistent with patterns observed across the full seasonal dataset (Fig. S7). These four focal modules differed in size and network topology (Fig. 6C). They were enriched for distinct biological processes relevant to stress acclimation and seasonal adjustment, including transport, RNA metabolism, photosynthesis, vascular development, and hormone-related signaling (Fig. 6D; Table S13).

**Figure 6.**
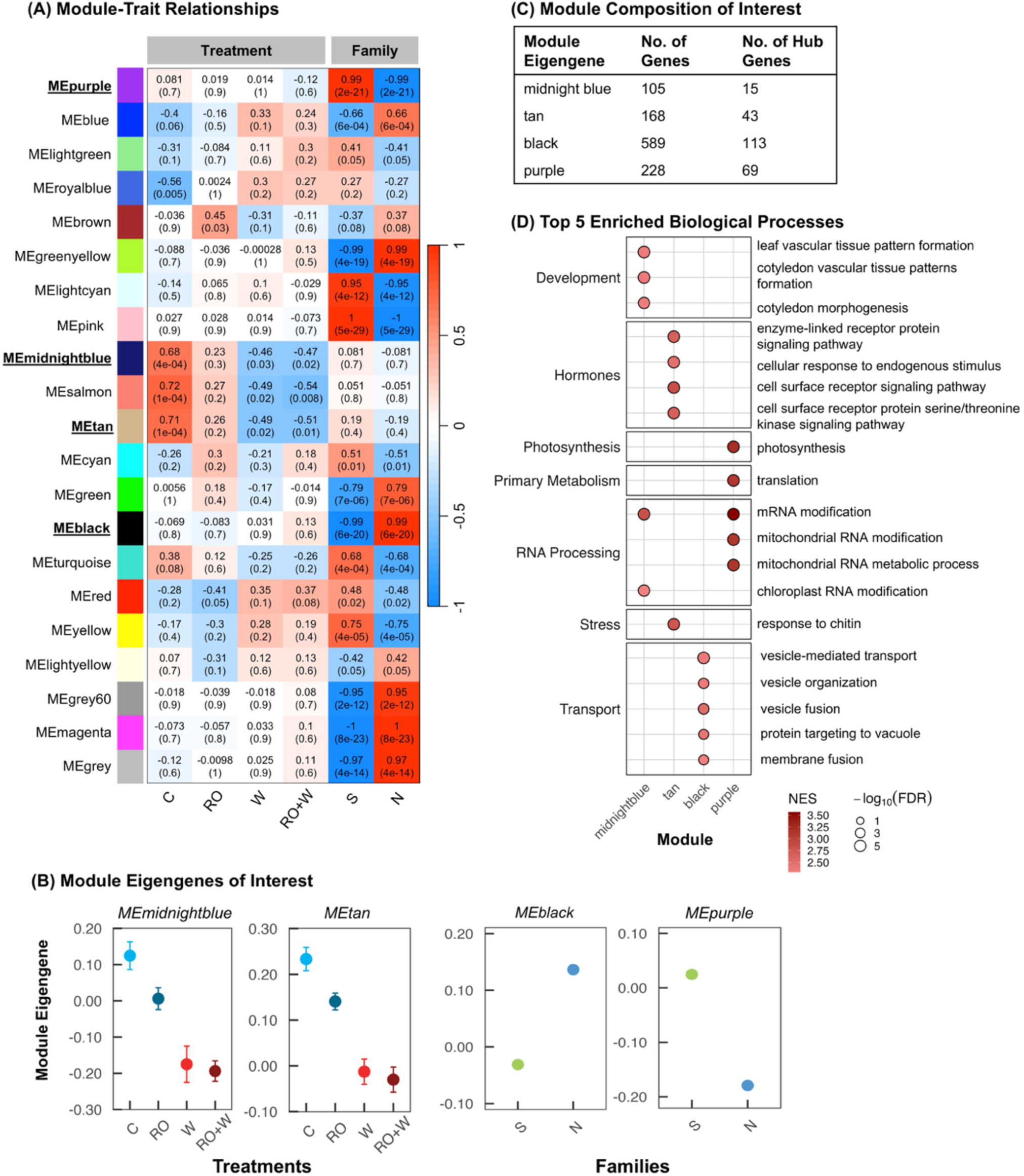
December-specific module–trait relationships, eigengene patterns, and functional enrichment of focal co-expression modules. (A) Module–trait heatmap showing Pearson correlations between December module eigengenes and treatment and family. Colours indicate correlation direction and strength (red, positive; blue, negative), with correlation coefficients and P-values shown. (B) Mean module eigengene values ± SE for focal modules across treatments (midnight blue, tan) and families (black, purple). (C) Summary of focal modules. (D) Top positively enriched GO Biological Process terms for each focal module based on GSEA. Dot colour represents normalized enrichment score (NES), and dot size represents −log10(FDR), capped at 5 (FDR=0.00001).

Annotated hub genes within treatment-associated modules closely reflected these functional themes and point to coordinated regulation of cold acclimation and stress signaling (Table S14). In the MEmidnightblue module, hub genes included homologs encoding transcription factors (*ZAT9, WRKY48*), and a calmodulin-binding protein kinase (*CaMRLK*). All have established roles in cold and oxidative stress responses. Several additional highly module-associated genes, including *CBP60f* and *LEA5* were also linked to cold acclimation. In the MEtan module, hub genes included homologs associated with drought and cold signaling, such as *PAL, COR47-like*, and *EIL1.* The module also contained *PYL12*, a key component of ABA signaling, which showed strong module association despite falling below the hub threshold. This is consistent with hormone-mediated stress signaling under late-season warming.

### Regulatory gene expression patterns across drought, cold, photoperiod, and light signalling pathways

We mapped differentially expressed transcripts (DETs) onto a published cold-acclimation signaling framework (Chang *et al*. 2021; Fig. 7). Putative white spruce homologs of regulatory genes associated with drought, cold, photoperiod, and light signaling were identified in the annotated long-read transcriptome and intersected with RNA-Seq results (Tables S15-19).

**Figure 7.**
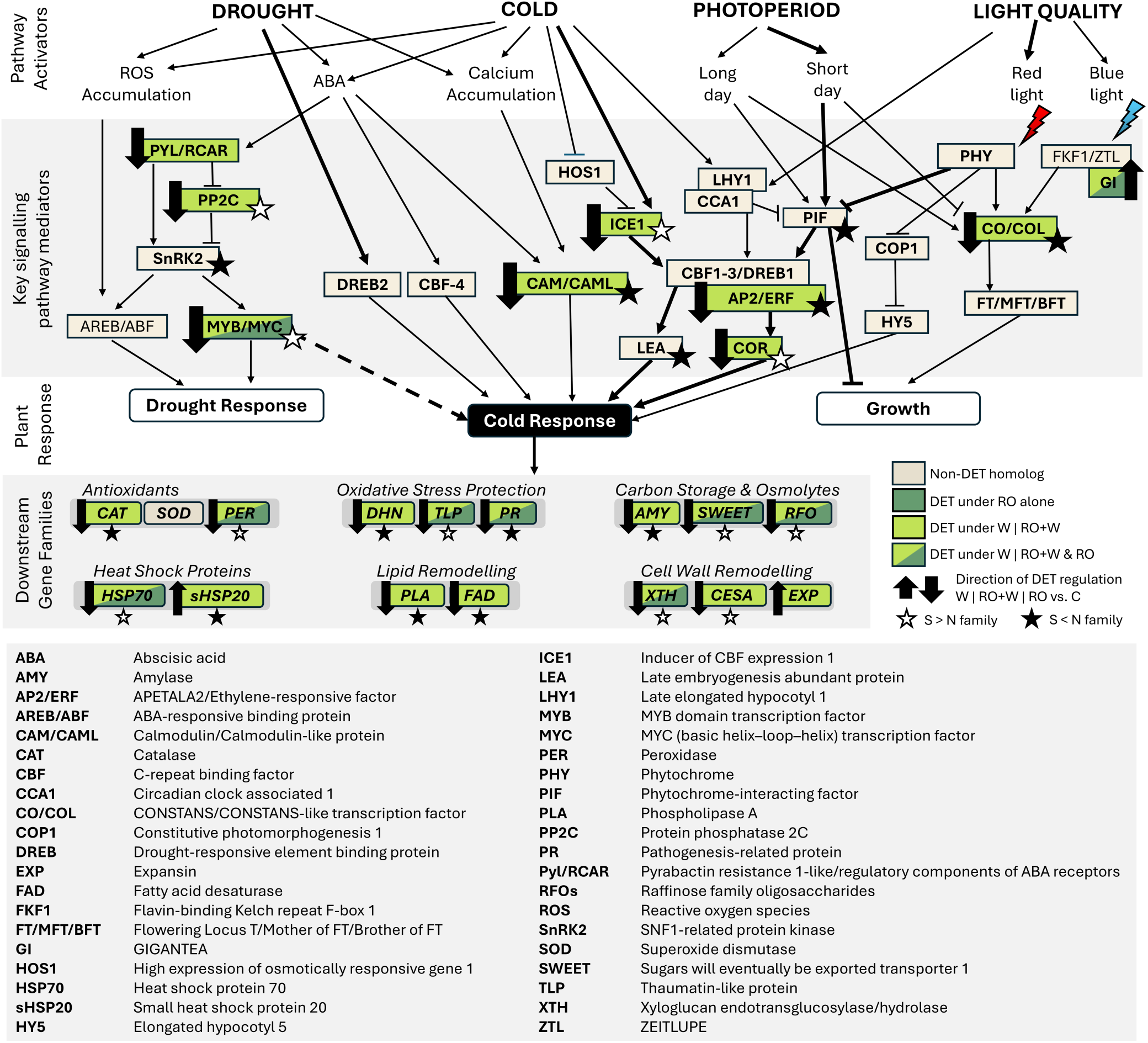
Summary of climate change impacts on the drought, cold, photoperiod and light signalling pathway of white spruce. Nodes represent pathway components and downstream gene families involved in stress signaling, growth regulation, and cold acclimation. Coloured boxes indicate detected homologs and differential expression status: light green denotes DETs under warming treatments (W or RO+W) relative to control, and dark green denotes DETs also observed under rainout (RO) alone. Differential expression is based on October and December contrasts only. Arrow direction indicates regulation under W or RO+W and, where applicable, was consistent with regulation observed under RO. Direction was determined from the mean log₂ fold change across homologous isoforms. Stars indicate significant Southern vs. Northern family differences within W or RO+W treatments. Arabidopsis-derived homologs are shown in plain font and conifer homologs in bold, with non-DETs included to provide pathway context. Central components of these networks are denoted by thick arrows. Figure represents composite pathways based on Chang *et al*. (2021).

Across pathways, several upstream signaling components were differentially expressed under W and RO+W. Abscisic acid–related regulators (*PYL/RCAR, PP2C*), along with downstream MYB/MYC transcription factors, were downregulated under W, with MYB/MYC transcript expression also reduced under RO. Cold-associated regulatory components, including *ICE1*, *COR*, CAM/CAM-like, and AP2/ERF family members, were likewise downregulated under W. Consistent with this pattern, downstream gene families involved in stress protection (PR, TLP), lipid remodeling (PLA, FAD), and carbohydrate metabolism (AMY, SWEET) were downregulated under both W and RO (Fig. 7). In contrast, photoperiod- and light-associated regulators (*LHY1, CCA1, PIF, PHY, COP1*) showed limited response to either treatments. Homologs of *CO/CO-like* genes were downregulated and *GI* was upregulated under W, whereas other downstream photoperiod-related regulators showed no detectable response.

Family-level differences within this regulatory framework varied across treatments. Most cold-response regulatory and downstream gene families showed lower transcript abundance in the southern family than in the northern family under both W and RO+W. The southern family also showed reduced expression of photoperiod- and light-signaling genes (*PIF, CO/CO-like*) under W and RO+W compared to the northern family (Fig. 7). Homologs of both *CO-like* and *FT-like* genes were downregulated in the southern family in July, even under control conditions (Table S17).

## Discussion

Cold acclimation is essential for evergreen conifers to survive freezing winters at northern latitudes and is a key determinant of boreal forest productivity under climate change. In the northern hemisphere, conifers are locally adapted to climatic gradients, with cold acclimation representing a major axis of selection. This adaptation reflects a fundamental trade-off between maximizing late-season carbon gain and ensuring timely acquisition of freezing tolerance. Consequently, forest health and productivity depend on the tight coordination between photosynthesis, growth cessation and cold acclimation under future climates. Warmer autumn conditions can extend photosynthetic activity later into the season (Chang *et al*., 2016; Fréchette *et al*., 2016; Stinziano & Way, 2017), increasing late-season carbon gain. However, warming can also delay the acquisition of freezing tolerance (Repo *et al*., 1996; Guak *et al*., 1998; Chang *et al*., 2016). This response is not universal, as some conifer species maintain similar or enhanced freezing tolerance under experimental warming (Kreyling *et al*., 2012; Noordermeer *et al*., 2021). Despite extensive work on phenological and physiological responses to late-season warming, the molecular regulation coordinating late-season photosynthesis with cold acclimation has received little attention. Here, we showed that autumn warming maintained photosynthesis-related regulatory programs while reducing expression of the transcriptional cascade associated with cold acclimation. The moderate summer drought imposed by the rainout (RO) treatment had limited effects across most physiological and transcriptomic responses.

### Autumn warming extends expression of photosynthetic genes whereas summer drought has minimal effects

Warming induced the largest treatment effects on late-season physiology and photosynthesis-related transcriptional patterns. Warming delayed photosynthetic downregulation into December, as indicated by higher values of PSII photochemistry and carotenoid-based vegetation indices (Murphy *et al*. 2025; Fig. S1). Expression of genes involved in carbon fixation, thylakoid maintenance, and electron transport remained elevated under warming as temperatures declined in the control treatment (Figs. 4 and S3; Table S9). Notably, this response included Calvin cycle enzymes and regulatory genes linked to photosystem efficiency (*CYP37;* Yang *et al*., 2023). Together, these patterns indicate that warmer autumn conditions prolong photosynthetic capacity into early winter rather than accelerating seasonal downregulation.

RO effects were limited across phenotypic traits (Murphy *et al. 2025;* Fig. S1), gene set enrichment and differential expression analysis, showing minimal carryover of moderate summer drought onto autumn transcriptome reprogramming. RO yielded the fewest differentially expressed transcripts and rarely altered the direction of warming responses (Figs. 3, 4 and S4). The RO+W treatment closely resembled W, indicating that temperature was the dominant climatic driver of late-season regulation in this experiment (Fig. 3C). Although photosynthesis-related gene sets were enriched under RO, few individual transcripts were differentially expressed. Gene set enrichment instead identified subtle but coordinated shifts in gene expression. The limited impact of RO likely reflects both its moderate severity and timing. Decreases in soil moisture were greatest during summer, before autumn cold acclimation intensified (Fig. S1A). More severe or later-season water limitation can accelerate photosynthetic downregulation in conifers (Wang *et al*., 2025). Late-season drought therefore remains a potential modifier of seasonal transitions under warming.

### Family identity shapes photosynthetic responses to warming

Family identity explained baseline differences in gene expression and shaped the timing and magnitude of warming responses. Principal component analysis reinforced this distinction. Family explained the largest share of variation (PC1, 32%), whereas warming defined a secondary axis (PC2, ∼11.5%) in October and December (Fig. S2). Gene set enrichment further revealed family-dependent responses to warming (Fig. 4). The southern family maintained higher photosynthetic activity later into autumn and showed higher expression of carbon fixation and photosystem-related genes under both W and RO+W (Fig. S4). Divergence between families was most pronounced for genes associated with PSII maintenance and chloroplast function, including *TL29*, *CLPB2*, and *THF1*, across treatments and time points (Fig. S5). Sustained expression of PSII and electron transport genes contributes to photosystem integrity and repair (Aro *et al*., 1993; Su *et al*., 2023). These results indicate persistent family-level differences in the maintenance of the photosynthetic machinery with consequences for carbon acquisition under warming.

### Warming suppresses cold acclimation regulatory networks independently of CBF pathways

Warming reduced enrichment of gene sets related to transcriptional regulation and signaling into early winter (Fig. 4). This shift coincided with the downregulation of multiple putative transcription factor families, driven predominantly by AP2/ERF and MYB superfamilies (Fig. S6). Although CBF/DREB1 transcription factors are central to herbaceous cold-response models (Fowler & Thomashow, 2002; Cook *et al*., 2004; Yamaguchi-Shinozaki & Shinozaki, 2006), warming had limited effects on the expression of *CBF-like* homologs. Instead, reductions occurred in upstream signaling components and downstream cold-response markers, including *ICE1*-, *CAM*-, and *COR*-like homologs (Figs. 5 and 7). This challenges the prevailing model derived largely from herbaceous systems and points to a broader regulatory architecture in conifers. Evidence from Norway spruce also shows that cold acclimation in conifers may be dominated by non-CBF regulatory programs involving AP2/ERF, MYB, and WRKY-like genes (Vergara *et al*., 2022). Our results suggest that AP2/ERF and MYB-associated regulation contributes more substantially to cold acclimation in conifers than models derived largely from herbaceous systems would predict (Chang *et al*., 2021).

These shifts in transcriptional regulation extended to the downregulation of multiple downstream gene families (Fig. 7; Table S18), indicating a reduced activation of key protective pathways normally engaged during cold acclimation. Genes associated with antioxidant metabolism (e.g., *NEC2, PER67-like*) and oxidative stress protection (e.g., *COR47-like*, *RPV1-like*) showed reduced expression, suggesting diminished capacity for ROS detoxification. Reduced expression of carbohydrate-related genes, including *GolS2*, involved in osmoprotective sugar synthesis, and *SWEET1*, a sugar transporter, suggests weaker osmotic adjustment. Decreased expression of heat shock proteins (*HSP70*) indicates less investment in protein stabilization under stress. Warming may therefore suppress the coordinated suite of molecular responses that prepare tissues for freezing tolerance (Steponkus, 1984; Guy, 1990; Strimbeck *et al*., 2008; Chang *et al*., 2021). The magnitude and direction of these effects on downstream genes also varied by family, indicating that warming effects on cold acclimation are genotype-dependent. Comparable provenance-linked variation in downstream cold acclimation gene families has been reported in Sitka spruce across latitudinal gradients (Holliday *et al*., 2008). Co-expression analysis further identified modules enriched for regulatory and stress-response functions that were downregulated under warming, indicating reduced coordination of transcriptional control, hormone signaling, and cold acclimation at the network level.

### Growth cessation is photoperiod-controlled and insensitive to warming in white spruce

These regulatory changes under warming occurred without detectable shifts in the timing of growth cessation. Under ambient conditions, gene sets related to growth cessation declined shortly after the summer solstice (Fig. 2C; Table S3). This timing aligns with established xylem phenology in boreal conifers, where cambial activity and growth cessation track daylength more closely than temperature (Repo *et al*., 2000; Rossi *et al*., 2006; Ježík *et al*., 2021). The southern family nonetheless showed reduced expression of several core clock and photoperiod regulators under control and warming, including *PIF-like* and *CO-like* genes. The FT expression also differed between families during July (Table S17). Latitudinal variation in *FT-like* gene regulation has been linked to the timing of growth cessation in Norway and Siberian spruce (Gyllenstrand *et al*., 2007; Li *et al*., 2019). In August under ambient conditions, the northern family also showed an eightfold greater number of differentially expressed transcripts compared to the southern family (Fig. 2A), likely reflecting earlier initiation of bud set as well as growth cessation (Gyllenstrand *et al*., 2007; Guo *et al*., 2022). Together, these early-season differences appear to reflect variation in photoperiod pathway activity between families, with implications for adaptive variation in growth phenology.

### Warming alters cell wall and xylem-associated gene expression in a family-dependent manner

Following growth cessation, seasonal regulation shifts toward late-season structural development, including latewood differentiation and secondary wall deposition. Latewood formation occurs within a broader acclimation window in late summer to early autumn (Cartenì *et al*., 2018). During this time, temperature affects xylem differentiation and latewood traits (Begum *et al*., 2012, 2018; Fonti *et al*., 2013; He *et al*., 2025). In our experiment, warming effects differed by family, particularly for hydraulic diameter and tracheid wall reinforcement (Fig. S1D; Murphy *et al*., 2025).

Cold acclimation in herbaceous systems involves coordinated remodeling of cell walls and membranes that reduces cellular injury during freezing (Steponkus, 1984; Yamada *et al*., 2002). This involves shifts in wall polysaccharide composition and turnover, induction of wall-modifying enzymes, and regulation of phenylpropanoid and lignin-related pathways (Le *et al*., 2015; Ji *et al*., 2015; Takahashi *et al*., 2019; Kutsuno *et al*., 2023). Here, homologs associated with cellulose synthesis (*CESA5)*, cell-wall remodeling (*XTH8*), and phenylpropanoid metabolism (*PAL-like*) showed reduced expression under warming treatments in both families (Fig. S4; Table S10). However, transcriptomic patterns by family did not align with the xylem anatomy. For example, the southern family downregulated *CCOAOMT*, a phenylpropanoid pathway gene linked to cell wall reinforcement, in October under W and RO+W (Fig. S5), despite increased latewood tracheid wall reinforcement and relative wood density observed under warming (Fig. S1D; Murphy *et al*. 2025). This mismatch likely reflects tissue differences between needle transcriptomes and stem xylem anatomy. Warming instead likely altered the timing and coordination of late-season wall-building processes in needle tissue, while family-dependent responses were more evident in stem xylem. Extended photosynthetic activity in the southern family under warming may have also increased sugar availability, promoting thicker latewood cell walls (Cartenì *et al*., 2018).

### Conclusions

Autumn warming sustained photosynthesis-associated transcriptional programs into early winter while suppressing cold-acclimation networks, demonstrating that temperature-sensitive processes can become uncoupled from photoperiod-regulated growth cessation in white spruce. Contrary to predictions from herbaceous model systems, this response was not strongly associated with canonical CBF/DREB regulators but instead involved MYB- and AP2/ERF-centered transcriptional networks, suggesting a distinct mechanism of cold acclimation in conifers. Together, these findings identify transcriptional mechanisms underlying previously observed warming-induced delays in photosynthetic downregulation and provide evidence that warmer autumns may weaken cold acclimation without extending growth in boreal conifers. By identifying the regulatory basis of this decoupling, our findings contribute to resolving the broader uncertainty surrounding the coupling between carbon uptake and growth in forests under climate change (Rao *et al*., 2026).

## Acknowledgements

We gratefully acknowledge the Canadian Forest Service Quebec, in particular Daniel Plourde and Eric Dussault, for their contributions to seedling production, planting, and technical support during experimental setup. We thank the Institute for Systems Genomics Computational Biology Core at the University of Connecticut (UConn) for access and assistance with computational resources. Generative AI (Copilot, GPT-5.5) was used to assist with code optimization and copy editing of the manuscript.

## Author Contributions

IE, NI, and BKM conceptualized the experiment. JW and VMEV guided the bioinformatics workflow. BKM performed experimental work. BKM and NP analyzed the data. BKM and IE wrote the manuscript. All authors have reviewed the manuscript.

## Data Availability

Long-read RNA-seq libraries and the associated *de novo* transcriptome assembly (GLNV00000000), along with short-read RNA-seq libraries generated from white spruce BioSamples (accession nos. SAMN60423273 to SAMN60423374), have been deposited in NCBI BioProject PRJNA1471275 and are currently under embargo pending publication of Murphy *et al*. (in press). Supporting datasets, analysis scripts, and other files necessary to reproduce the analyses are archived and available to reviewers upon request. Public access will be provided upon publication.

## Supplemental Information

### Supplemental Methods

- **Methods S1**. Long-read *de novo* transcriptome assembly and annotation pipeline from Murphy et al. (press).
- **Method S2.** Treatment implementation and environmental monitoring.

### Supplemental Figures

- **Figure S1.** Experimental design and selected results from autumn cold acclimation experiment.
- **Figure S2.** Principal component analysis (PCA) of normalized gene expression data for October and December time points.
- **Figure S3.** Broad functional biological GO pathway enrichment across climate treatments and family contrasts.
- **Figure S4.** Candidate photosynthesis- and xylem-related transcripts showing repeated differential expression in the control versus applied treatment contrasts.
- **Figure S5.** Candidate photosynthesis- and xylem-related transcripts showing repeated differential expression in the southern versus northern family.
- **Figure S6.** Distribution of differentially expressed transcription factors among transcription factor families in October (A–C) and December (D–F).
- **Figure S7.** Module–trait relationships across the full dataset (all months).

### Supplemental Tables

- **Table S1.** Summary of bioinformatics output for Nova-Seq short-read sequencing libraries generated from mature needles of white spruce seedlings.
- **Table S2.** Differentially expressed genes identified from two white spruce *(Picea glauca)* families of contrasting provenance (northern and southern) under ambient (control) conditions comparing July to August, October and December.
- **Table S3.** Gene Ontology (GO) Biological Process enrichment results from gene set enrichment analyses (fgsea) comparing July to later seasonal sampling months (August, October, and December) within each family under ambient (control) conditions only.
- **Table S4.** Differentially expressed genes identified from two white spruce (*Picea glauca*) families of contrasting provenance (northern and southern) comparing control vs. applied treatments (RO, W, RO+W) in August, October and December.
- **Table S5.** Overlap of treatment-responsive differentially expressed transcripts (DETs) within families in October and December.
- **Table S6.** Gene Ontology (GO) gene set enrichment results from fgsea analyses across all treatment × family comparisons.
- **Table S7.** Most consistently enriched up- and GO Biological, Molecular, and Cellular Process terms across treatment and family contrasts.
- **Table S8.** Gene Ontology (GO) and KEGG terms used to define biological pathways of interest for pathway-based candidate gene identification.
- **Table S9**. Ranked candidate transcripts associated with photosynthesis and related carbon metabolism pathways.
- **Table S10.** Ranked candidate transcripts associated with xylem- and cell-wall biogenesis metabolism pathways.
- **Table S11.** Differentially expressed transcription factor transcripts identified across sampling months, provenance families, and treatments.
- **Table S12.** The 50 most variable transcription factors across sampling months and treatments, including transcript identifiers, transcription factor family assignments, average expression values (AveExpr), and predicted *Arabidopsis thaliana* homologs.
- **Table S13.** All significantly enriched GO Biological Process gene sets for all co-expression modules. This table reports significant gene set enrichment results for each co-expression module.
- **Table S14.** Top hub genes per co-expression module based on intramodular connectivity.
- **Table S15**. Similarity search results for candidate cold acclimation regulators.
- **Table S16**. *De novo* transcriptome annotation matches for transcript identifiers recovered in Table S15.
- **Table S17.** Differential expression summary for candidate homologs identified in Tables S15 and S16.
- **Table S18.** Downstream gene family homologs and differential expression directionality for W and RO+W contrasts only.
- **Table S19.** Downstream gene family homologs and differential expression directionality for RO contrasts only.

