## Supplemental Materials and Figures for "Autumn warming decouples the molecular control of photosynthesis and cold acclimation from growth cessation in white spruce"

Bridget K. Murphy, Noelle Perkins, Vera M.E. Velasco, Nathalie Isabel,

Jill Wegrzyn, and Ingo Ensminger

**Supplemental Methods S1. Long-read *de novo* transcriptome assembly and annotation pipeline from Murphy et al. (accepted).**

For long-read (LR) sequencing, one biological replicate per family, treatment, and time point was selected from the Autumn Cold Acclimation Experiment (n = 22). One biological replicate per family was also selected from the control treatment at the August time point of a Summer RO and Warming Experiment (n = 2). In total, 24 LR-RNA libraries were generated. LR-RNA sequencing was performed at The Centre for Applied Genomics (TCAG) in Toronto, Canada. For each sample, 500 ng of total RNA was processed using the Iso‑Seq Express Library Preparation workflow with the SMRTbell Express Template Prep Kit 2.0 (Pacific Biosciences, Menlo Park, CA, USA), following the manufacturer’s protocol. Briefly, full‑length barcoded cDNA was synthesized, amplified, and pooled equimolarly into groups of six samples. Each pool was converted into a SMRTbell library and sequenced on a single SMRT 8M cell using the PacBio Sequel IIe platform with a 24‑hour movie time.

Iso‑Seq LR-RNA sequencing data generated on the PacBio Sequel II platform were processed using the Iso‑Seq3 Analysis Workflow implemented in SMRT Link. Subreads were demultiplexed by barcode, followed by removal of 5′ and 3′ cDNA primers. Reads shorter than 50 bp were excluded, and poly(A) tails and concatemers were removed to generate full‑length non‑chimeric (FLNC) reads. FLNC reads were clustered and iteratively merged to produce unpolished transcript models, which were subsequently polished to improve consensus accuracy and generate high‑quality transcripts used for downstream analyses. Coding sequence (CDS) regions were predicted from high‑quality Iso‑Seq transcripts using TransDecoder v5.5.0 (https://github.com/TransDecoder). Within each Iso‑Seq library, predicted CDS regions were clustered at 95% sequence identity using VSEARCH v2.4.3 (Rognes *et al.*, 2016) to remove redundancy and generate sets of unique transcripts. Unique CDS regions from all 24 libraries were then clustered at 80% sequence identity to produce a pre‑filtered *de novo* LR transcriptome prior to contaminant removal.

Functional gene annotation and contamination filtering were performed using EnTAP v2.3.0 (Hart *et al.*, 2020). Sequence similarity searches within EnTAP were conducted using DIAMOND against the Plant Protein Database release 232, UniProt (v2025.06), Araport11, and a custom gymnosperm protein database comprising 186,061 sequences from seven species (*Picea abies, Picea sitchensis, Ginkgo biloba, Cycas micholitzii, Gnetum montanum, Taxus baccata*, and *Abies sachalinensis*; available at <https://gitlab.com/PlantGenomicsLab/HQ_Douglas-fir_transcriptome_genome_annotation>).

Gene Ontology (GO) annotations within EnTAP were assigned using eggNOG‑mapper v2.0.1 (Cantalapiedra *et al.*, 2021) with eggNOG v5.0.2, UniProt v2025.06, and InterProScan v5.7.6 databases. Transcripts whose best‑supported taxonomic assignment corresponded to Bacteria, Fungi, Insecta, or Mammalia were classified as contaminants and removed. Transcriptome assembly quality was assessed using rnaQUAST (Bushmanova *et al.*, 2016) and BUSCO v5.4.5 (Simão *et al.*, 2015; Manni *et al.*, 2021) with the Embryophyta lineage dataset from OrthoDB. The LR *de novo* transcriptome demonstrated high assembly quality, with 93.1% BUSCO Embryophyta completeness and functional annotation assigned to 65% of transcripts, supporting its use as a reference for RNA-seq analyses.

**Supplementary Method S2. Treatment implementation and environmental monitoring**

Rainout structures were installed between seedlings using longitudinally halved PVC profiles placed at an incline to intercept and divert approximately 50% of precipitation away from the rooting zone (Fig S1E). Non-rainout plots contained dummy profiles oriented to allow rainfall to drip into the soil to control for structural effects. Rainout structures were removed on November 27 in preparation for snowfall. Warming was implemented using an infrared heater array in a T-FACE configuration (Fréchette *et al.*, 2020) operated from August 31 through late December to maintain an approximately +5 °C increase in leaf temperature relative to ambient conditions.

Each plot was equipped with sensors for air temperature and humidity, soil temperature and moisture, and infrared leaf temperature, logged at 30-min intervals using a CR1000 datalogger (Campbell Scientific). An on-site weather station recorded air temperature and humidity, PAR, solar flux, and precipitation at 15-min intervals. Occasional gaps in precipitation measurements were filled using records from the King City North station (Environment Canada), located 8.5 km from the field site.

**Supplemental Figures**

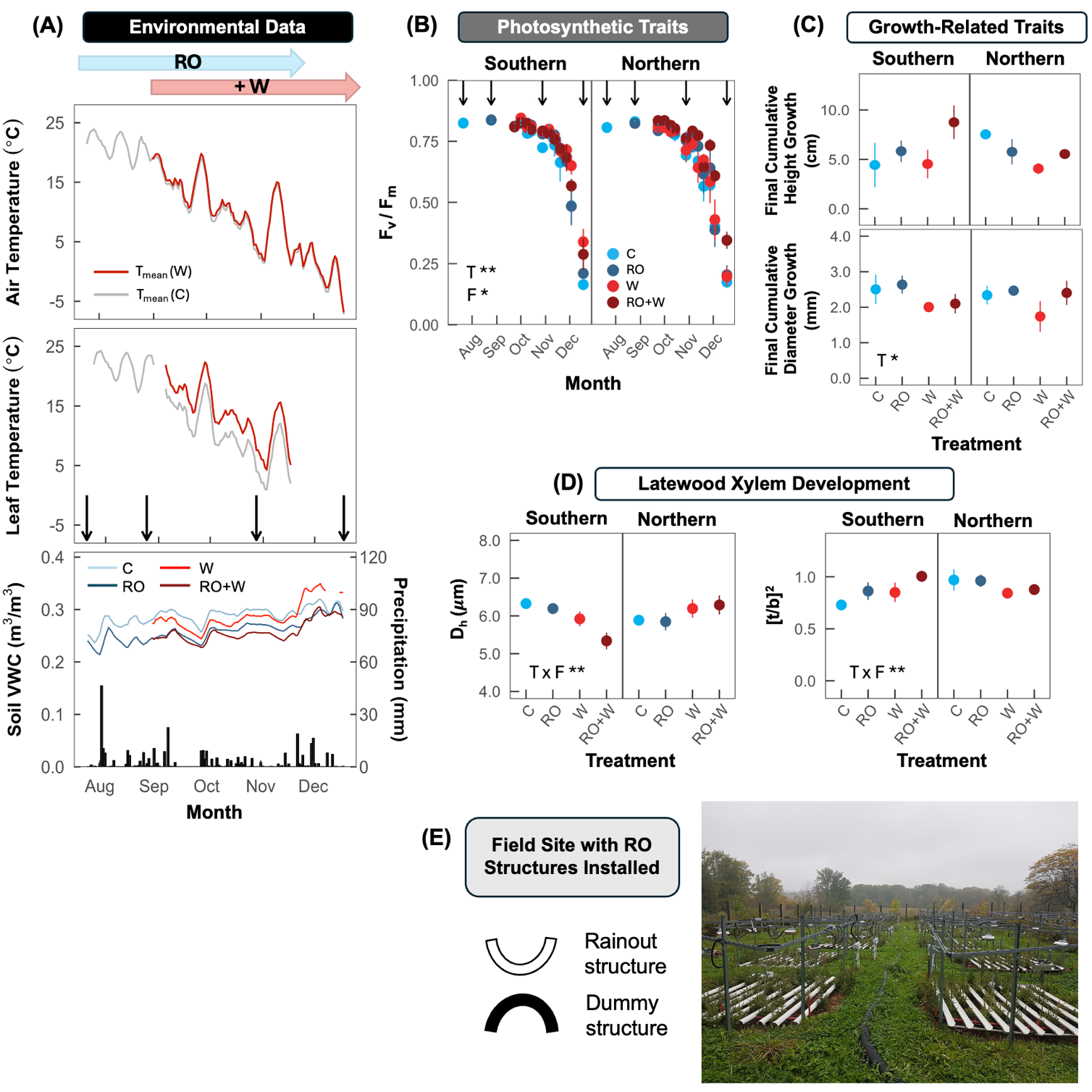

**Supplemental Figure S1. Experimental design and selected results from autumn cold acclimation experiment.** (A) Environmental conditions, including air temperature, leaf temperature, and soil volumetric water content (VWC); (B) One of four measured photosynthetic traits, maximum PSII efficiency (F_v_/F_m_); (C) Final cumulative height and diameter growth; (D) Two of six measured latewood xylem anatomical traits, hydraulic diameter (D_h_) and tracheid wall reinforcement, [t/b]². (E) Photo of field site and a schematic of the rainout and dummy structures. C, control; RO, rainout; W, warming; RO+W, warming plus rainout. Points are plot means, error bars are ± SE (n = 3–12). Fixed effects were temperature (T), soil water content (SWC), and family (F); only significant main effects or interactions are shown (P < 0.05). *P ≤ 0.05, **P < 0.01, ***P < 0.001. Black arrows in panels (A) and (B) indicate sampling dates for gene expression. Adapted from Murphy *et al.* (2025).

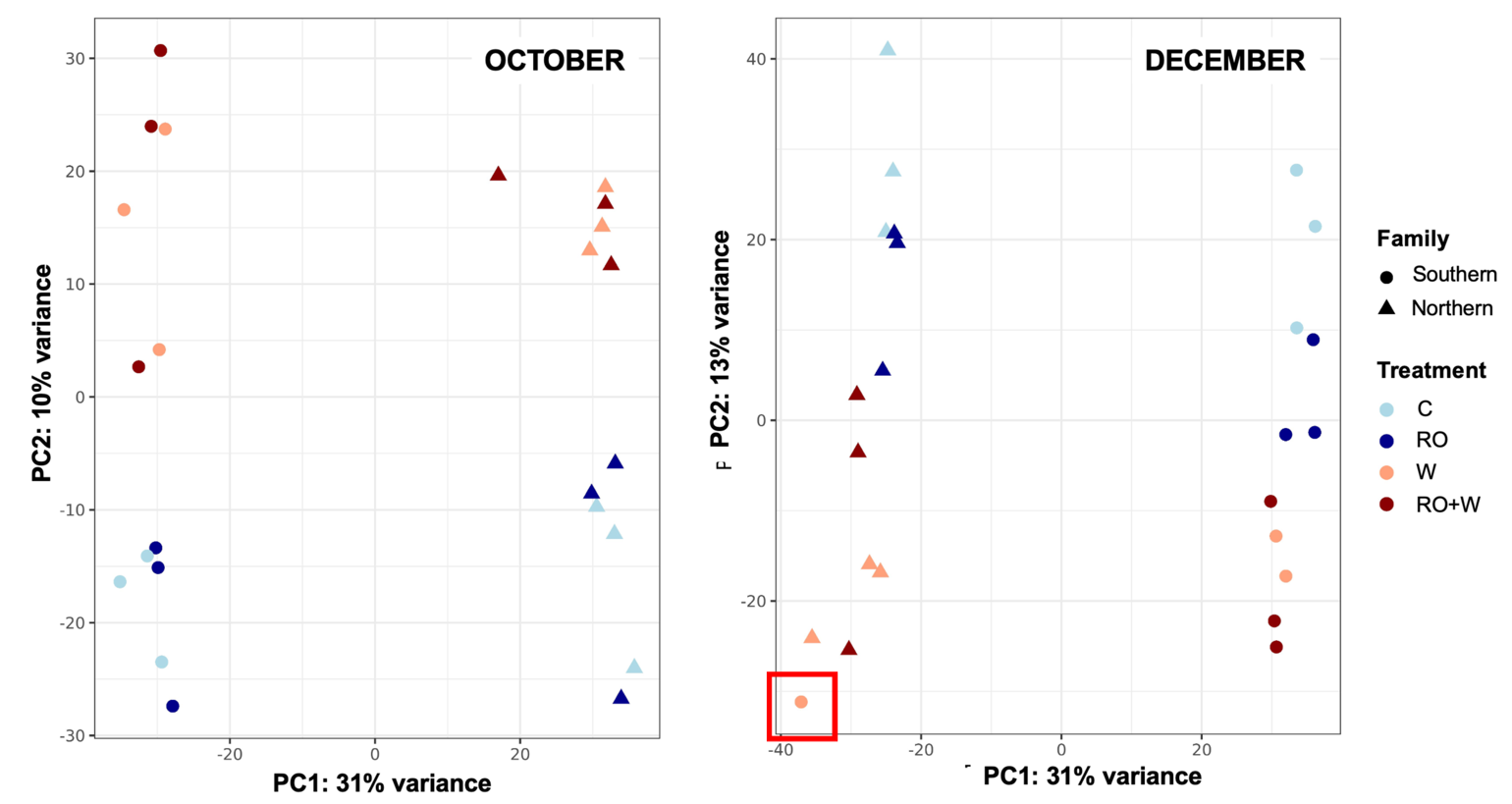

**Supplemental Figure S2. Principal component analysis (PCA) of normalized gene expression data for October and December time points.** PCA was performed on DESeq2-normalized counts from all samples to visualize overall expression patterns and treatment effects. Points represent individual samples, colored by treatment (C, RO, W, RO+W) and shaped by family (Northern, Southern). The mislabelled sample from the December subset is indicated with a red square outline.

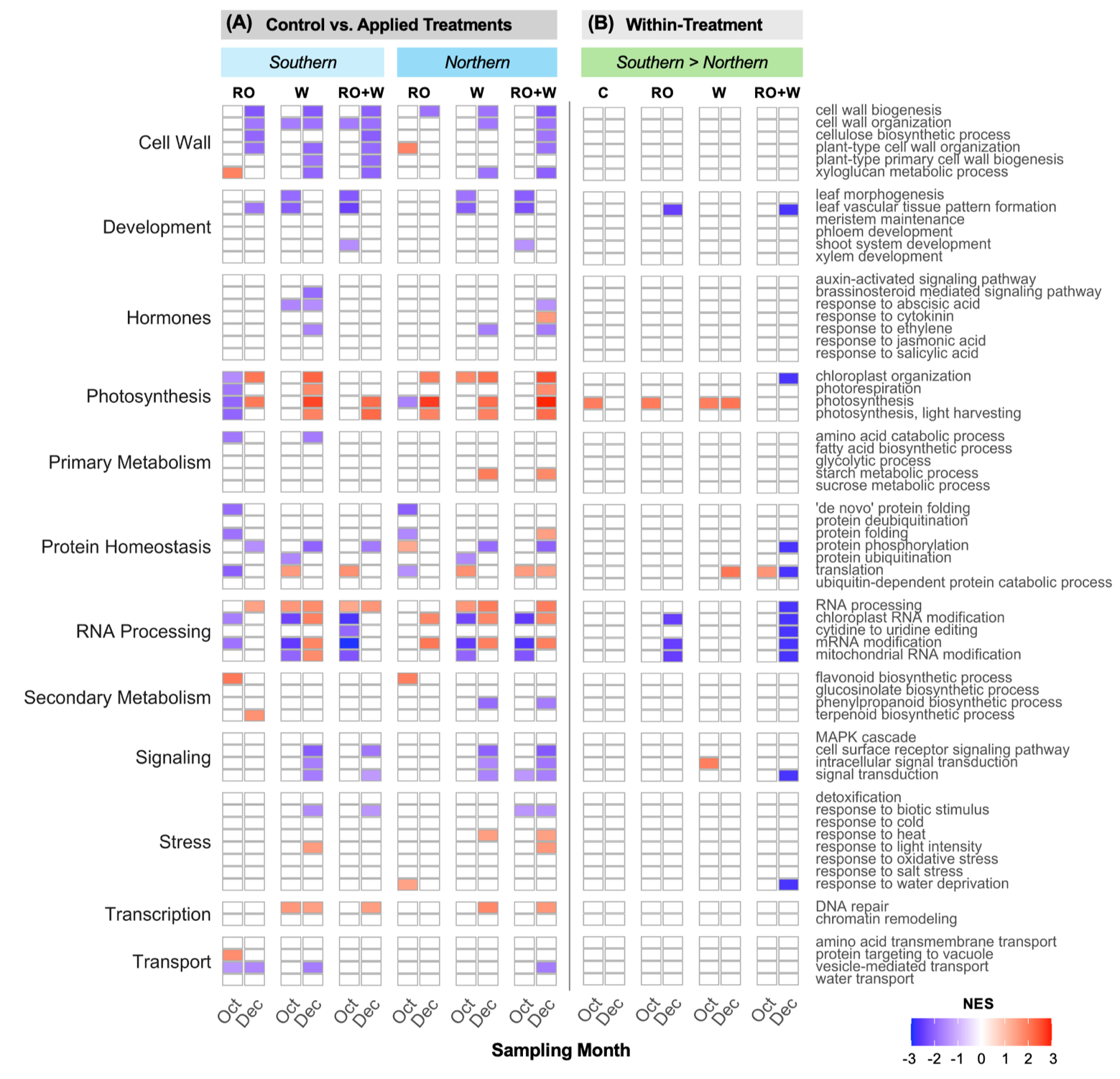

**Supplemental Figure S3. Broad functional biological GO pathway enrichment across climate treatments and family contrasts.** Heatmap shows normalized enrichment scores (NES) from gene set enrichment analysis (GSEA) for **(A)** treatment contrasts: C versus RO, W, and RO+W within each family; and **(B)** family contrasts within treatments, comparing Southern versus Northern within each treatment, including the control. Results are shown for October and December. Tiles are coloured by NES only when enrichment was significant (FDR < 0.05), with red indicating positive NES and blue indicating negative NES. For Control vs. Applied Treatments, positive NES indicates enrichment toward transcripts higher in the applied treatment relative to control. For Southern > Northern, positive NES indicates enrichment toward transcripts higher in the southern family relative to the northern family. Uncoloured cells indicate non-significant enrichment. GO terms are grouped by functional category. C, control; RO, rainout; W, warming; RO+W, rainout combined with warming.

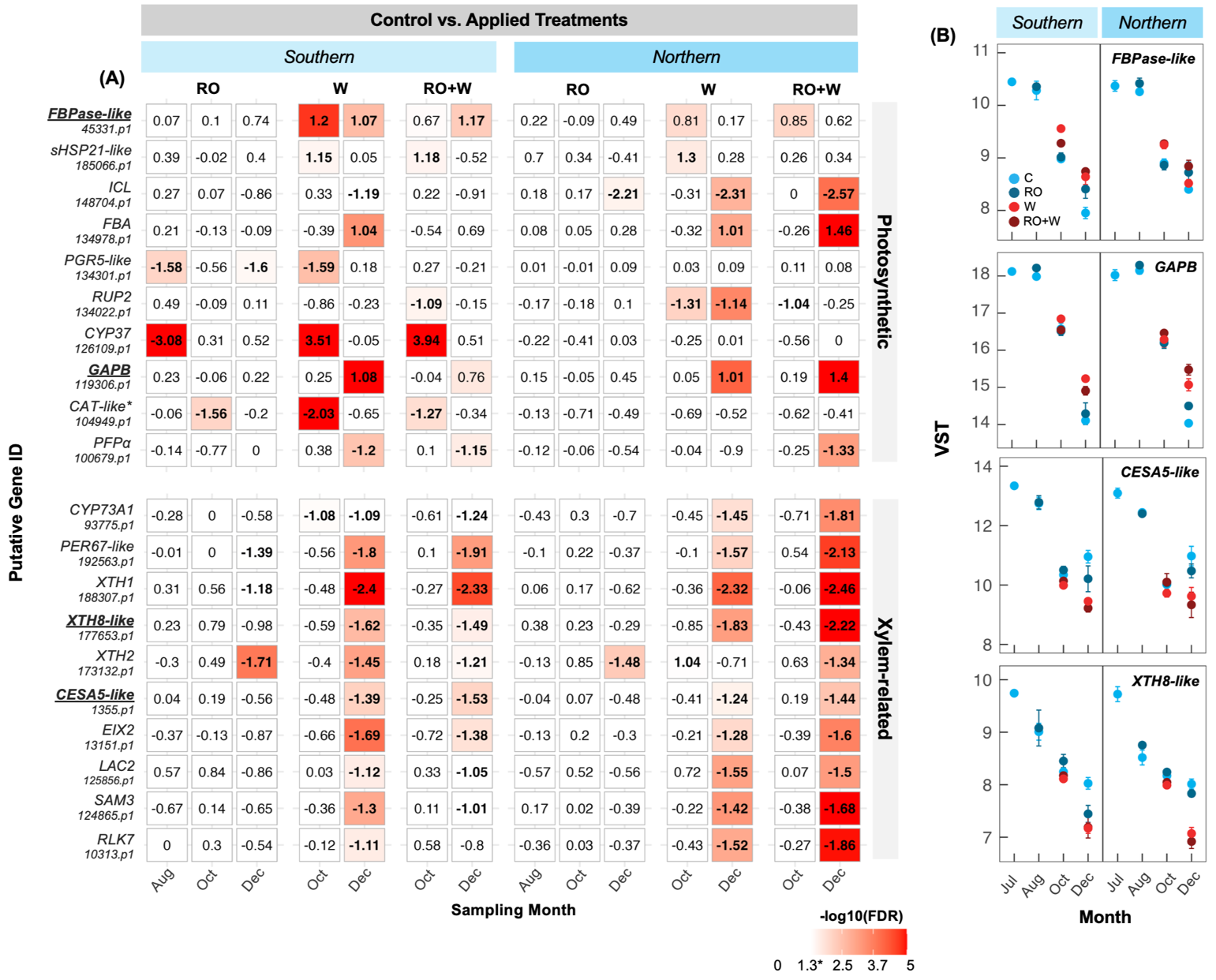

**Supplemental Figure S4. Candidate photosynthesis‑ and xylem‑related transcripts showing repeated differential expression in the control versus applied treatment contrasts.** (A) Heatmap of control versus applied treatment contrasts (RO, W, RO+W) within Southern and Northern families, evaluated independently at each sampling month (August, October, December). Cell colour denotes significance (−log10[FDR]), capped at 5 (FDR=0.00001) with 1.3 corresponding to FDR=0.05. Values indicate log2 fold change (LFC), with bold type showing |LFC| > 1. Transcripts are grouped by functional category based on GO and KEGG annotations. (B) Variance stabilizing transformation (VST) gene expression values across sampling months (July-December) for selected candidate transcripts, shown separately for Southern and Northern families and coloured by treatment. GO and KEGG terms defining pathway groupings are listed in Table S8. C, control; RO, rainout; W, warming; RO+W, combined rainout and warming. Gene names marked with an asterisk indicate annotations derived from gene family assignment only.

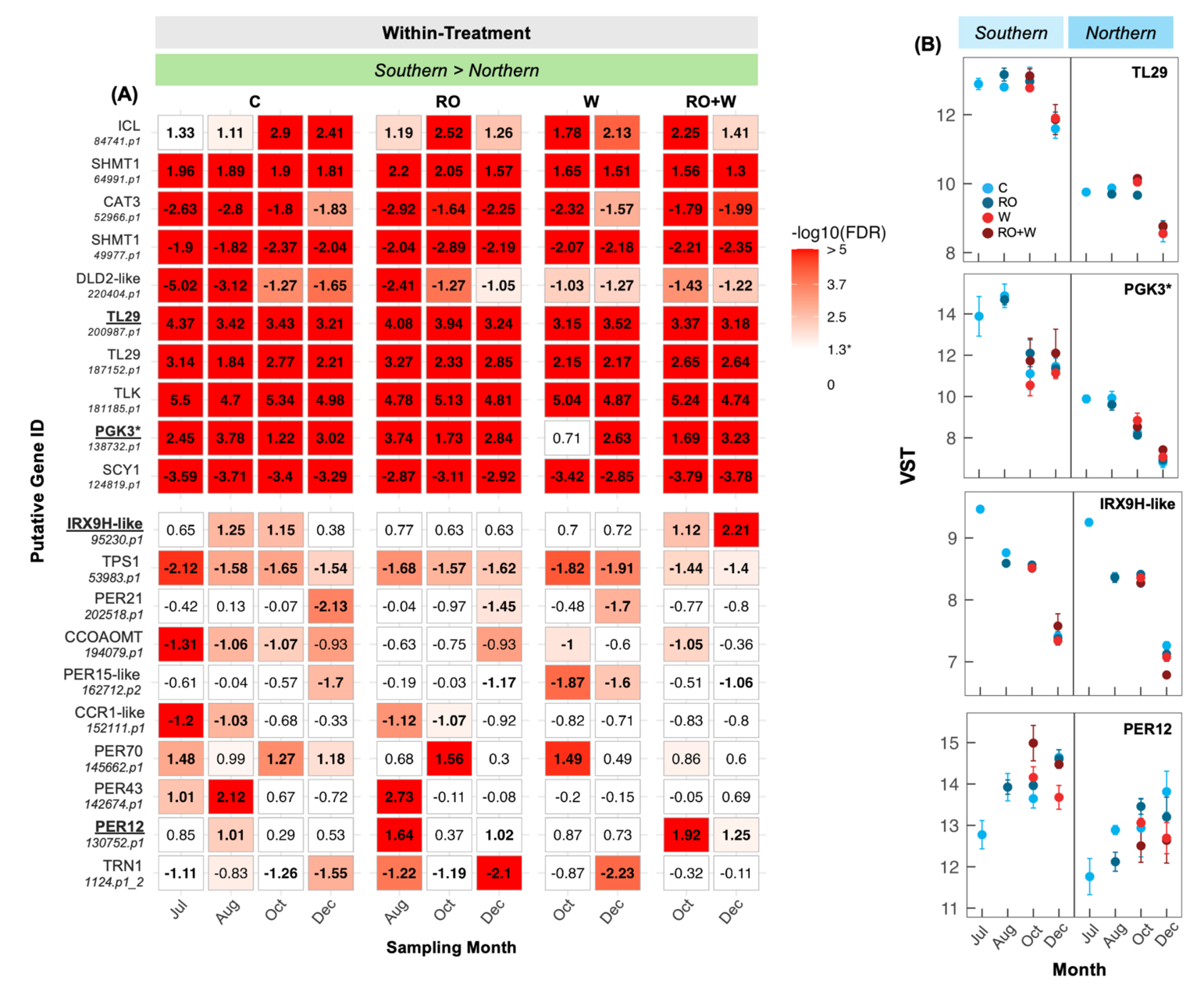
**Supplemental Figure S5. Candidate photosynthesis‑ and xylem‑related transcripts showing repeated differential expression in the southern versus northern family.** (A) Heatmap of within-treatment family contrasts (Southern > Northern) under each treatment, including the control, evaluated independently at each sampling month (August, October, December). Cell colour denotes significance (−log10[FDR]), capped at 5. Values indicate log2 fold change (LFC), with bold type showing |LFC| > 1. Transcripts are grouped by functional category based on GO and KEGG annotations. (B) DESeq2‑normalized counts across sampling months (July–December) for selected candidate transcripts, shown separately for Southern and Northern families and coloured by treatment. GO and KEGG terms defining pathway groupings are listed in Supplemental Table S8. C, control; RO, rainout; W, warming; RO+W, combined rainout and warming. Gene names marked with an asterisk indicate annotations derived from gene family assignment only.

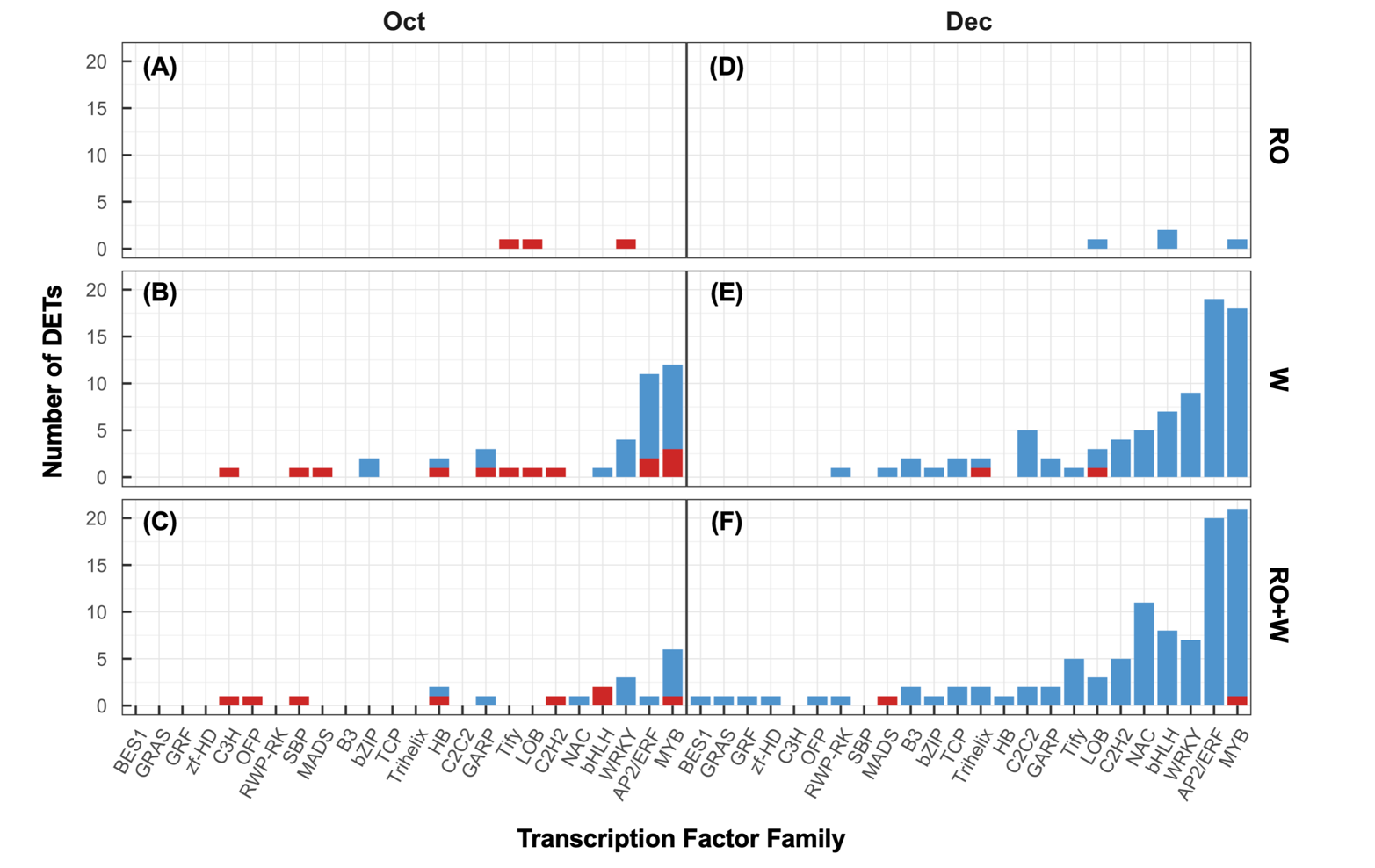
**Supplemental Figure S6. Distribution of differentially expressed transcription factors among transcription factor families in October (A–C) and December (D–F).** Panels represent RO (A, D), W (B, E), and RO+W (C, F) treatments relative to C. Bars indicate the number of transcription factors differentially expressed within each family, separated by direction of expression change (upregulated in red, downregulated in blue). C, control; RO, rainout; W, warming; RO+W, rainout combined with warming. Transcription factor families were assigned based on PlantTFDB classifications, with closely related subfamilies pooled into broader family categories (e.g., AP2/ERF, MYB, NF‑Y) for visualization and summary analyses.

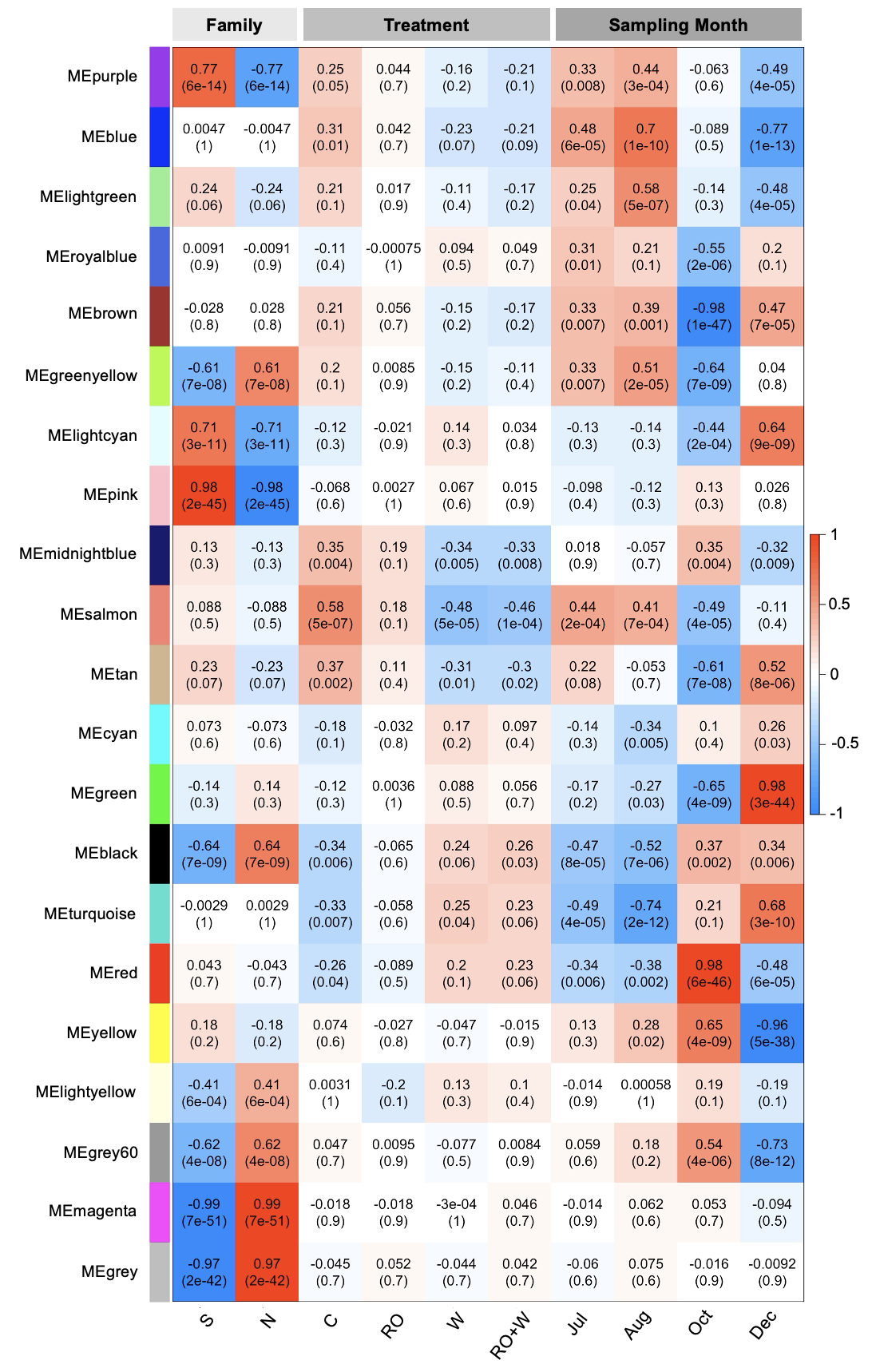

**Supplemental Figure S7. Module–trait relationships across the full dataset (all months).** Heatmap shows Pearson correlations between module eigengenes (rows) and experimental and temporal variables (columns), including family (S, N), treatment (C, RO, W, RO+W), and sampling month (Jul, Aug, Oct, Dec). Cell colours indicate the direction and strength of correlations (red, positive; blue, negative), and values in each cell report the correlation coefficient with the corresponding P‑value in parentheses. Modules are labeled by colour (for example, MEpurple, MEblue), and the colour key indicates correlation values ranging from −1 to 1.

**Supplemental Table Legends**

*[Attached as an excel file with separate sheets per supplemental table and a table of contents]*

**Table S1. Summary of bioinformatics output for Nova-Seq short-read sequencing libraries generated from mature needles of white spruce seedlings.** Number of raw reads, trimmed reads, assembled reads, identified coding regions, and unique transcripts are shown. Data shows mean ± SE for each family by treatment. Families are as follows: 437, Hungerford ON [Southern]; 495, Lac Simard, QE [Northern].

**Table S2. Differentially expressed genes identified from two white spruce (*Picea glauca*) families of contrasting provenance (northern and southern) under ambient (control) conditions comparing July to August, October and December.** Transcripts were mapped to the white spruce de novo transcriptome using Kallisto. DESeq2 was applied with an absolute log2foldchange (LFC) threshold of 1 and an adjusted p-value cutoff of 0.05. The transcriptome was annotated using EnTAP.

**Table S3. Gene Ontology (GO) Biological Process enrichment results from gene set enrichment analyses (fgsea) comparing July to later seasonal sampling months (August, October, and December) within each family under ambient (control) conditions only.** Reported statistics include GO identifiers and term names, normalized enrichment scores (NES), multiple‑testing–corrected P‑values, leading‑edge genes, and derived measures of enrichment direction. The target sampling month for each comparison is included. Only enrichment results with P_adj_ <0.05 are included in the table.

**Table S4. Differentially expressed genes identified from two white spruce (Picea glauca) families of contrasting provenance (northern and southern) comparing control vs. applied treatments (RO, W, RO+W) in August, October and December.** Transcripts were mapped to the white spruce *de novo* transcriptome using Kallisto. DESeq2 was applied with an absolute log2foldchange (LFC) threshold of 1 and an adjusted p-value cutoff of 0.05. The transcriptome was annotated using EnTAP.

**Table S5. Overlap of treatment-responsive differentially expressed transcripts (DETs) within families in October and December.** The table lists transcripts contributing to each overlap category shown in the Venn diagram (Fig. 3C), including unique and shared DETs among treatment contrasts (RO, W, and RO+W) within each family in October and December. For each transcript, family, sampling month, and overlap category are indicated, along with sequence annotations. Transcript annotations represent putative white spruce homologs derived from sequence similarity and functional databases.

**Table S6. Gene Ontology (GO) gene set enrichment results from fgsea analyses across all treatment × family comparisons**. The table reports GO identifiers and term names, normalized enrichment scores (NES), P‑values, FDR‑adjusted P‑values, gene set sizes, and leading‑edge genes, with comparison details. Only enrichment results with P_adj_ <0.05 are included in the table.

**Table S7. Most consistently enriched up- and GO Biological, Molecular, and Cellular Process terms across treatment and family contrasts.** The five most recurrent GO Biological, Molecular and Cellular Process terms in the up‑ and down‑regulated directions are shown for treatment‑level (control vs treatment within family) and family‑level (southern vs northern within treatment) comparisons. Listed contrasts indicate the pairwise fgsea comparisons included in each summary. Rankings prioritize the number of significant contrasts (nsignificant), with mean normalized enrichment score (NES) used to resolve ties. nₜₒₜₐₗ reflects the total number of available contrasts across months. GO terms associated with animal‑specific biology were excluded prior to analysis.

**Table S8. Gene Ontology (GO) and KEGG terms used to define biological pathways of interest for pathway‑based candidate gene identification.** Terms are grouped by biological category reflecting core processes, associated structural functions, or regulatory mechanisms relevant to the pathways analysed in this study.

**Table S9. Ranked candidate transcripts associated with photosynthesis and related carbon metabolism pathways.** Candidates were required to be significant in multiple contrasts spanning sampling month, family, and treatment. Within this pathway group, transcripts were ranked by the number of significant contrasts (n_combinations.x), with ties broken by the typical absolute log2 fold change (typical_abs_LFC). Functional annotation fields shown in this table are reported from the de novo long‑read transcriptome annotations. Both applied treatment contrasts within families (C vs RO, C vs W, C vs RO+W) and within‑treatment family contrasts (Southern > Northern) are included (contrast_type), and the contributing contrast combinations are listed in combinations with the associated sampling month.

**Table S10. Ranked candidate transcripts associated with xylem- and cell-wall biogenesis metabolism pathways.** Candidates were required to be significant in multiple contrasts spanning sampling month, family, and treatment. Within this pathway group, transcripts were ranked by the number of significant contrasts (n_combinations.x), with ties broken by the typical absolute log2 fold change (typical_abs_LFC). Functional annotation fields shown in this table are reported from the de novo long‑read transcriptome annotations. Both applied treatment contrasts within families (C vs RO, C vs W, C vs RO+W) and within‑treatment family contrasts (Southern > Northern) are included (contrast_type), and the contributing contrast combinations are listed in combinations with the associated sampling month.

**Table S11. Differentially expressed transcription factor transcripts identified across sampling months, provenance families, and treatments.** Reported values include transcript identifiers, transcription factor family assignment, log₂ fold change, adjusted p‑value (p_adj_), sampling month, provenance family, treatment, and direction of differential expression relative to the corresponding control.

**Table S12. The 50 most variable transcription factors across sampling months and treatments, including transcript identifiers, transcription factor family assignments, average expression values (AveExpr), and predicted *Arabidopsis thaliana* homologs.** Sequence similarity metrics (percent identity, mismatches, gap openings, alignment coordinates, and E‑values) and subject annotations are reported for each transcript.

**Table S13. All significantly enriched GO Biological Process gene sets for all co-expression modules. This table reports significant gene set enrichment results for each co-expression module.** Columns are: pathway (gene set identifier), p_val_ (nominal enrichment P‑value), p_adj_ (FDR‑adjusted P‑value), log2err (standard error for enrichment score estimates), ES (enrichment score), NES (normalized enrichment score), size (gene set size), leadingEdge (leading-edge subset of genes driving the enrichment signal), Module (module assignment), and GO_name (gene set description).

**Table S14. Top hub genes per co‑expression module based on intramodular connectivity. For each module, the top hub genes are listed, ranked by HubScore, a combined hub metric calculated as kWithin × MM, where kWithin is intramodular connectivity and MM is module membership (correlation with the module eigengene).** The table reports gene identifier (Gene), module assignment (Module), intramodular connectivity (kWithin), module membership (MM), and the combined hub metric (HubScore). Functional annotation fields shown in this table are reported from the *de novo* long‑read transcriptome annotations. Hub genes were computed from the full co-expression network, using the module assignments defined from the full dataset.

**Table S15. Similarity search results for candidate cold acclimation regulators.** Similarity search outputs from EnTAP v2.3.0 annotation of the long-read de novo white spruce transcriptome, used to identify putative homologs of genes included in the cold acclimation pathway framework (3). Results are shown for searches against Araport11 and RefSeq plant protein databases and include alignment and annotation fields for each query.

**Table S16. *De novo* transcriptome annotation matches for transcript identifiers recovered in Table S15.** Long-read *de novo* transcriptome annotations corresponding to the transcript identifiers identified as putative homologs in Table S15. Entries provide the matched annotation records for each unique transcript identifier, facilitating downstream cross-referencing with differential expression results.

**Table S17. Differential expression summary for candidate homologs identified in Tables S15 and S16.** RNA-Seq differential expression results for unique transcript identifiers corresponding to candidate cold acclimation regulators identified by similarity search (Table S15) and confirmed through *de novo* annotations (Table S16). Table reports statistical significance and log₂ fold-change estimates under warming, reduced soil water content, and combined treatments relative to control conditions.

**Table S18. Downstream gene family homologs and differential expression directionality for W and RO+W contrasts only.** Gene-family level summaries of annotated homologs from EnTAP similarity searches, linked to RNA-Seq contrasts. Includes dominant log₂ fold-change direction, direction-consistency metrics, counts of positive and negative responses, minimum FDR, and supporting functional annotation fields (GO, KEGG, UniProt, EggNOG, InterProScan). Limited to Oct/Dec sampling points.

**Table S19. Downstream gene family homologs and differential expression directionality for RO contrasts only.** Gene-family level summaries of annotated homologs from EnTAP similarity searches, linked to RNA-Seq contrasts. Includes dominant log₂ fold-change direction, direction-consistency metrics, counts of positive and negative responses, minimum FDR, and supporting functional annotation fields (GO, KEGG, UniProt, EggNOG, InterProScan). Limited to Oct/Dec sampling points
